# Flexibly Modeling Rare Variant Pathogenicity Improves Gene Discovery for Complex Traits

**DOI:** 10.1101/2025.11.04.686320

**Authors:** Jeremy Schwartzentruber, Petko P. Fiziev, Jeremy McRae, Jacob C. Ulirsch, Kyle Kai-How Farh

## Abstract

Rare variant burden tests can directly identify genes that influence complex traits, but their power is limited by our ability to separate functional from benign alleles. We introduce FlexRV, an approach that greatly improves the power to detect gene-based associations in rare variant aggregation tests by modelling nonlinear relationships between functional annotations and phenotype. Across 62 quantitative and 44 disease traits in the UK Biobank, we show that FlexRV outperforms previous approaches such as DeepRVAT, STAAR, and Regenie, discovering 51% more quantitative and 102% more disease trait associations than the widely used Regenie method. Compared to discoveries from other methods, gene-phenotype associations identified by FlexRV replicated at a higher rate in the independent All of Us cohort and were more highly enriched at genes nominated by common variant genome-wide association studies. We explore the genetic architecture of complex traits using FlexRV burden tests, finding nearly equal contributions from missense and loss of function variants to rare variant burden heritability. FlexRV weights can also be incorporated into rare variant polygenic scores, improving their ability to identify individuals with extreme phenotypes. Our study illustrates the benefits of modelling nonlinear relationships between annotated variant effects and their downstream phenotypes in rare variant studies.

## Introduction

A major goal in genetics is to identify the protein-coding genes connected to human phenotypes. Genome-wide association studies (GWASs) of common variants have identified tens of thousands of associations with human traits and diseases^1^. However, it remains difficult to confidently link these associations to specific genes since most associated loci are in non-coding regions of the genome and each locus frequently contains multiple associated variants in linkage disequilibrium (LD)^2–4^. In contrast, rare variants typically have low LD with neighboring variants and are more abundant^5,6^. While tests of individual rare variants are more limited in power due to lower allele counts, aggregating these variants into functional groups — such as protein-altering variants within a gene — can enable well-powered assessments of their joint impact on a phenotype.

To date, most gene-based discovery efforts have selected variants to use in these aggregation tests by applying hard thresholds for predicted pathogenicity and allele frequency^7–9^. However, creating powerful variant “masks” relies not only on accurate prediction of a variant’s effect, which remains an incompletely solved problem, but selection of appropriate thresholds, which are unknown *a priori.* To address this uncertainty, previous efforts generated and tested multiple masks^7–9^. Although some power is lost after accounting for an increased number of tests, modern statistical approaches such as the Cauchy combination test (CCT)^10^ maintain calibration but permit more gene-trait discoveries than overly conservative approaches (*e.g.* Bonferroni)^11–14^.

One major limitation of variant masks is that they assume all included variants have the same effect size on the trait or disease of interest, which is unlikely in practice. To address this, most rare variant tests allow instead for each variant’s effect to be scaled by a quantitative weight, but how to choose optimal weights remains an open question^15^.

Previous work often assigns higher weights for lower allele frequency variants, reflecting an assumption that these variants have larger effect sizes on average. Other approaches use functional annotations (*e.g.* CADD^16^, PrimateAI-3D^17^) to derive sets of variant weights, similar to the use of multiple masks^14,18^.

A recently introduced method, DeepRVAT^19^, trained a neural network to integrate multiple functional annotations into a predicted “gene impairment” weight, but this approach assumes that the mapping from variant annotations to pathogenicity (“gene impairment”) is not only the same across genes but the mapping from pathogenicity to trait is linear. Whether these assumptions are valid is unknown and has received relatively little attention. There are reasons to expect that nonlinear relationships will be widespread^20^, and such relationships have been reported in humans^21^.

Here, we introduce FlexRV, a general approach to rare variant association testing that allows for flexible, nonlinear relationships between variant annotations and traits of interest. FlexRV requires no training data and can be incorporated into standard rare variant association tests, such as burden and sequence kernel association tests (SKAT)^22^, across multiple implementations, including STAAR^14^, SAIGE-GENE+^13^, and Regenie^23^. Using only two annotations, PrimateAI-3D^10^ and allele frequency, to derive weights, FlexRV discovers more genes associated with both complex human traits and diseases than previous approaches. Discoveries by FlexRV were more enriched at GWAS loci and more likely to replicate in the independent All of Us cohort. We further find that the best performing FlexRV weights differ based on gene properties such as constraint, and for most genes are not those with a linear relationship between predicted pathogenicity and trait measurement.

## Results

### Overview and performance of FlexRV

To improve discovery in rare variant association tests, we developed FlexRV, which allows for nonlinear relationships between variant annotations and a phenotype of interest. FlexRV takes one or more variant annotations, such as pathogenicity scores, and minor allele frequency (MAF) or minor allele count (MAC) as input, performs a small number of transformations to obtain weights, and performs rare variant association testing using each weight vector, accounting for multiple testing using CCT^10^. Building on previous work^7^, we considered loss-of-function (LoF) and missense variants, assigning an initial “pathogenicity” of 1 to frameshift, stop gained, and canonical splice variants. We used PrimateAI-3D^17^ as the predicted pathogenicity score for missense variants based on its strong performance^7,24^ and our observation that FlexRV enhances power for all pathogenicity scores tested (section “Optimal burden weights vary according to gene properties”). We then performed 16 transformations of the pathogenicity score and 12 transformations of MAF, taking the product of these transformations to generate 192 sets of weights for each gene (**Fig. 1a**, **Extended Data Fig. 1**, **Methods**). Transformations were chosen to encode biologically plausible relationships. We expect that the magnitude of variant effects on a trait may increase with predicted pathogenicity, but this relationship may not be linear. To allow for nonlinearity, we selected beta distributions, which allow varying slopes, and cubic root functions, which allow smooth transition points. Transformations of MAF were simpler, including hard thresholds and beta distributions, which are commonly used.

**Fig. 1:**
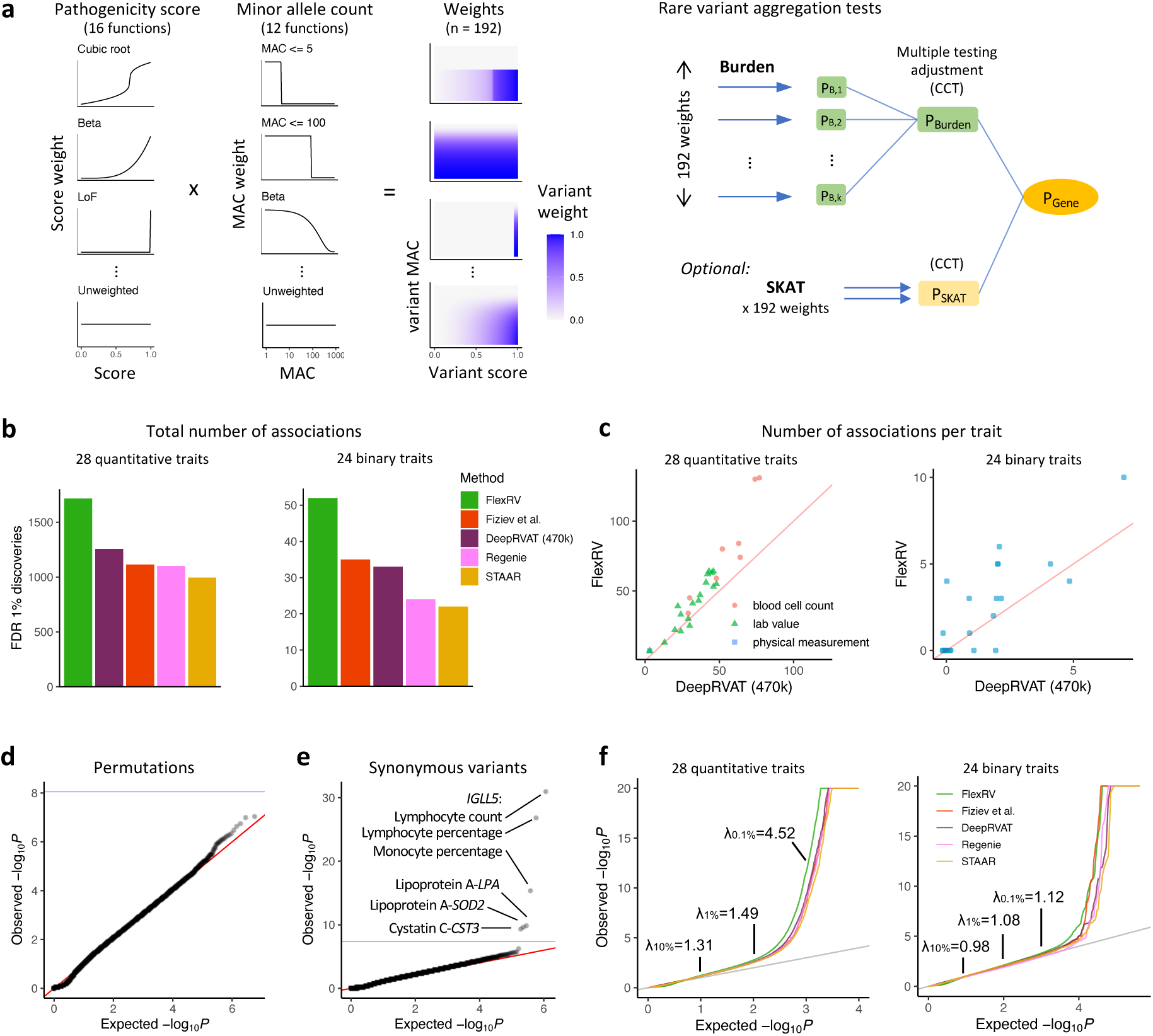
Overview and performance of FlexRV method. **a,** Variants are annotated with minor allele count (MAC) or frequency (MAF) and a variant pathogenicity score for missense variants (PrimateAI-3D). We define 16 transformations for variant score and 12 transformations for MAC. Their product defines 192 functions transforming variant annotations to weights. Each set of weights is used to test for association of variants in the gene with a phenotype (using Burden, SKAT, or other kernels). We applied FlexRV and alternative methods to 62 quantitative traits and 44 binary traits in the UK Biobank. **b**, Total number of gene-phenotype associations discovered by each method, across 28 quantitative and 24 binary traits. **c**, Scatterplot of the number of associations discovered by FlexRV vs. DeepRVAT for each trait. DeepRVAT had a slightly larger sample size based on the full 470k exome cohort. **d**, Quantile-quantile (QQ) plot of observed vs. expected p-values from burden tests where phenotype data was permuted, based on 5 permutations for each of 62 quantitative traits. **e**, QQ plot from burden tests using synonymous variants with real phenotypes for 62 quantitative traits, with significant trait-gene associations labelled. In both panels the blue line represents a Bonferroni-based significance threshold (0.05 / number of tests). **f**, True phenotype tests: QQ plot of observed vs. expected p-values. Inflation values (λ) of observed test statistics at different null p-value quantiles (top 10%, 1%, 0.1%) are shown.

Although FlexRV is general, we implement it using a generalised linear mixed model (GLMM) framework that handles covariates, population structure, and relatedness as in previous methods^14,23^. FlexRV supports all weighted kernel association tests^25^, including SKAT, although in this work we focus mainly on burden test results to facilitate comparisons across methods (Regenie and STAAR support SKAT tests, DeepRVAT and Fiziev et al.^7^ do not). Moreover, weights in Burden tests are more interpretable, since burden test power is optimized by weights that match true variant effect sizes, whereas optimal weights are less clear for SKAT. After performing each test, we obtained an omnibus p-value corrected for multiple testing using CCT.

Using 423,614 unrelated individuals of all ancestries in the UK Biobank (UKB) 450k whole exome release, we selected missense and LoF variants with MAF < 0.1% in 18,790 protein-coding genes. We performed weighted burden tests for 62 quantitative traits and 44 diseases (**Supplementary Table 1**) adjusting for age, sex, age x sex, UKB assessment center, 20 common variant genetic principal components (PCs), 20 rare variant genetic PCs, and a trait-specific common variant polygenic score (**Methods**).

We selected 4 alternative methods to compare against: Regenie^23^, STAAR^14^, DeepRVAT^19^, and our previous adaptive thresholding approach^7^. We ran Regenie, STAAR, and the Fiziev et al. approach on the exact same set of variants and traits as FlexRV. For Regenie, we used 4 MAF threshold masks and 3 annotation masks (LoF, LoF and missense, LoF and predicted deleterious missense), similar to Backman et al.^9^. For STAAR, we used variant pathogenicity scores for 10 annotation PCs from the FAVOR database^26^. To enable straightforward comparisons, we used CCT to obtain an omnibus p-value for both Regenie and STAAR. For Fiziev et al., we used the same pathogenicity and MAF annotations as FlexRV. For DeepRVAT, we considered published results on a set of 28 quantitative and 24 disease traits in a slightly larger subset of the UKB (∼470k whole exome release) that overlapped our analyses.

At a false discovery rate (FDR) of 1%, FlexRV discovered 2,807 burden test associations with quantitative traits and 123 gene-disease associations, outperforming all other methods (**Fig. 1b**, **Extended Data Fig. 2**). FlexRV identified 51% more quantitative trait associations and just over twice the number of binary disease trait associations as Regenie. Among genes identified by both, FlexRV typically had a smaller p-value (1257 / 1636 associations, 77%, **Extended Data Fig. 3**). Despite DeepRVAT having a slightly larger sample, FlexRV discovered 37% more quantitative and 58% more disease associations, with consistent improvements across traits (**Fig. 1b,c**). We also compared FlexRV burden, FlexRV SKAT, and FlexRV burden+SKAT tests (**Methods**), finding slightly fewer associations for SKAT (1,756) than burden, but 10% more for the combined burden and SKAT tests (3,103). Discoveries from all methods are reported in **Supplementary Tables 2-3**.

Although hypothesis tests from all methods are based on similar GLMMs and combined using the same previously validated CCT approach^10,14,27^, we sought to evaluate calibration of p-values reported by our method. We permuted phenotype residuals from a null model including only the covariates and re-ran FlexRV tests on quantitative and binary phenotypes. We observed slight inflation in test statistics in the range of p-values from 10^-2^ to 10^-4^ (average observed / expected 𝜒^2^ of 1.02 – 1.04) but inflation was negligible for p < 10⁻⁴, consistent with previous investigations into CCT^10,27^ (**Fig. 1d**). Across 5.7 million permuted gene-trait tests, none reached significance at an FDR of 1%. Despite control of false positives, we observed a strong deflation in median test statistics (average λ_gc_ = 0.22), suggesting that this metric may not be a reliable indicator of calibration for omnibus CCT tests, and the full spectrum of association results should be examined.

To further evaluate potential false positives, we ran FlexRV using only rare synonymous variants, which we expect to have little impact on traits (**Methods**). At an FDR of 1%, we found an association between *IGLL5* synonymous variants and three blood cell count traits (**Fig. 1e**), consistent with prior findings^8^. We also observed an association between lipoprotein A levels and synonymous variants in both *LPA* and *SOD2* (at the *LPA* locus). *LPA* harbors a variable number tandem repeat (VNTR) whose length has an extremely large effect on lipoprotein A levels^28^, which is likely in partial LD with these synonymous variants. Lastly, we found one association between *CST3* synonymous variants and cystatin C, which could similarly be explained by strong coding variant and CNV associations that were not fully accounted for^29^.

Returning to our primary results using LoF and missense variants, we examined quantile-quantile plots of observed and expected p-values (**Fig. 1f**). FlexRV and all alternative methods showed signal of non-null gene-trait associations beginning at expected p-values of 0.05 - 0.001 for quantitative traits, most notably for height and blood cell counts (**Extended Data Fig. 4**). To estimate whether this could be attributed to residual confounding or simply indicated a polygenic architecture, we used burden heritability regression (BHR)^30^. Intercepts estimated by the BHR model had a similar range to previous reports^30^, indicating minimal confounding (**Extended Data Fig. 5**).

### Validation and replication of discoveries

To evaluate replication of FlexRV results, we performed burden tests for 62 quantitative traits in a downsampled set of ∼200,000 unrelated individuals from UKB and investigated how many were significant in the larger UKB cohort, using results from the independent DeepRVAT method (Bonferroni p-value < 0.05, **Methods**). Compared to Regenie and STAAR, FlexRV discoveries replicated at a substantially higher rate across a range of significance ranks (**Fig. 2a**). We also sought to evaluate replication of FlexRV discoveries from the larger UKB cohort in the external All of Us (AoU) dataset^31^ for 18 matching quantitative traits which had at least one significant association in AoU (FDR < 1%). Using a more lenient significance threshold for replication in the smaller AoU cohort (p-value < 0.05), FlexRV discoveries replicated at an equal or greater rate than other methods, including DeepRVAT (**Fig. 2b**).

**Fig. 2:**
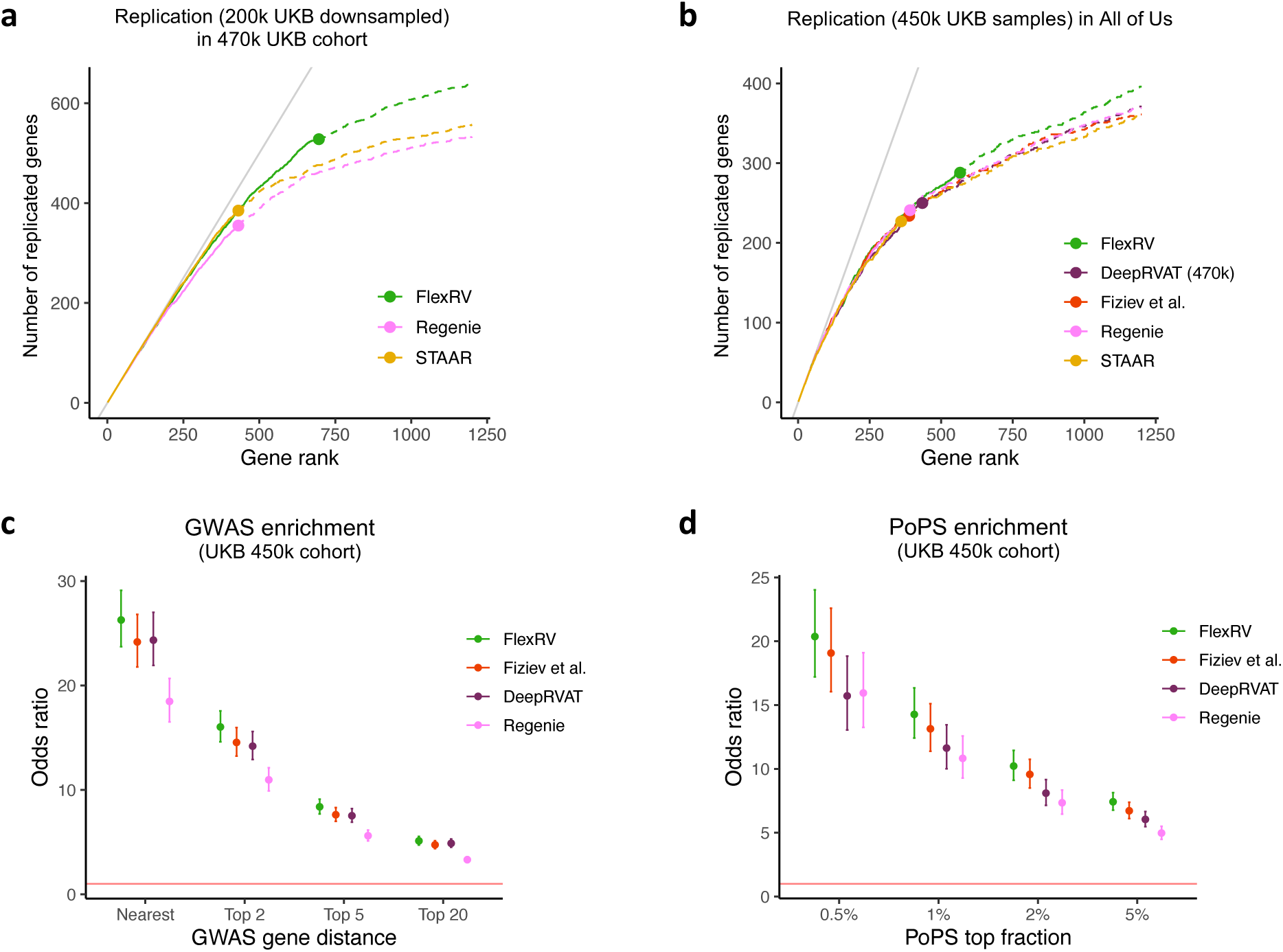
Validation and replication of discoveries. **a,** Number of top-N gene-trait associations discovered by each method (FlexRV, STAAR, Regenie) in a downsampled dataset of 200k UK Biobank individuals, which are replicated at nominal p-value < 2.5 x 10^-6^ in the larger cohort of 470k individuals as reported in Clarke et al.^12^, across 28 quantitative traits. The filled point for each method indicates the number of associations declared significant at FDR 1%, after which a dotted line shows replication of non-significant genes. **b,** Similar to (**a**), showing the number of top-N gene-trait associations discovered by each method in the 450k cohort which are replicated at p-value < 0.05 in the independent All Of Us cohort for a set of 18 overlapping quantitative traits. **c,** GWAS enrichment across 28 quantitative traits: considering the top 2,500 gene-trait associations by p-value from each method, shown is the enrichment of burden test genes for being the nearest, or among the nearest 2, 5, or 20 genes to a GWAS index SNP for the trait. **d,** For the same traits, we show the enrichment of the top 2,500 burden test associations from each method for having PoPS scores in the top percentiles across genes genome-wide for the matched trait. Error bars represent 95% CIs. The red lines in (**c**) and (**d**) indicate an odds ratio of 1.

As a final evaluation of FlexRV discoveries, we investigated whether they were supported by discoveries from common variant GWASs for the same quantitative traits in the same cohort. Although GWASs do not directly identify the trait-relevant causal genes, they are typically among the nearest genes to GWAS index variants^2,3^. We selected the top 2,500 genes by burden test p-value, approximately the number of FlexRV discoveries at FDR < 1%, and investigated if they were enriched among the genes close to a GWAS index variant. Each method was significantly enriched across multiple gene distance ranks, but FlexRV exhibited higher enrichment than DeepRVAT, Regenie, STAAR, and Fiziev et al. (**Fig. 2c**). An alternative way of identifying candidate causal genes from GWAS is PoPS^3^, a machine learning method which uses locus-independent features, such as cell type-specific expression, biological pathways, and protein-protein interactions, to rank trait-relevant genes not only at each GWAS locus but across the genome^32^. We evaluate FlexRV discoveries at top ranked PoPS genes, which are independent of specific GWAS loci, again finding that FlexRV is more significantly and more highly enriched than alternative methods (**Fig. 2d**).

### Optimal burden weights vary according to gene properties

Across 11 additional pathogenicity predictors, we found that FlexRV improved gene discovery in each, with an average improvement of 30% (**Extended Data Fig. 6,7**). To better understand how FlexRV achieves this performance, we visualized the relationship between the mean phenotype values of variant carriers and the variant pathogenicity score. We observed that this relationship was often nonlinear (**Fig. 3a,b**). After transforming the pathogenicity score to the best performing FlexRV burden weight, the phenotypic relationships were often more linear. For example, at *ASGR1*, the best-performing burden weight captures the transition from variants with little effect on alkaline phosphatase levels below a PrimateAI-3D score of ∼0.5 to those having a strong effect above ∼0.7 (**Fig. 3a**). At *TUBB1*, platelet count was similar for variant carriers with a PrimateAI-3D score below ∼0.7, but mean platelet count rapidly increased above this threshold (**Fig. 3b**). These trends remained consistent when other variant pathogenicity scores (*e.g.* AlphaMissense^33^, CADD^16^, REVEL^34^) were used instead of PrimateAI-3D (**Extended Data Fig. 8**) and were supported by many examples where FlexRV improved power with a nonlinear transformation (**Extended Data Fig. 9**). Indeed, LoF-only weights were the best-performing for only 4.5% of significant associations, and yet the best-performing transformations tended to be those with transition points at relatively high pathogenicity score values (**Fig. 3c,d**).

**Fig. 3:**
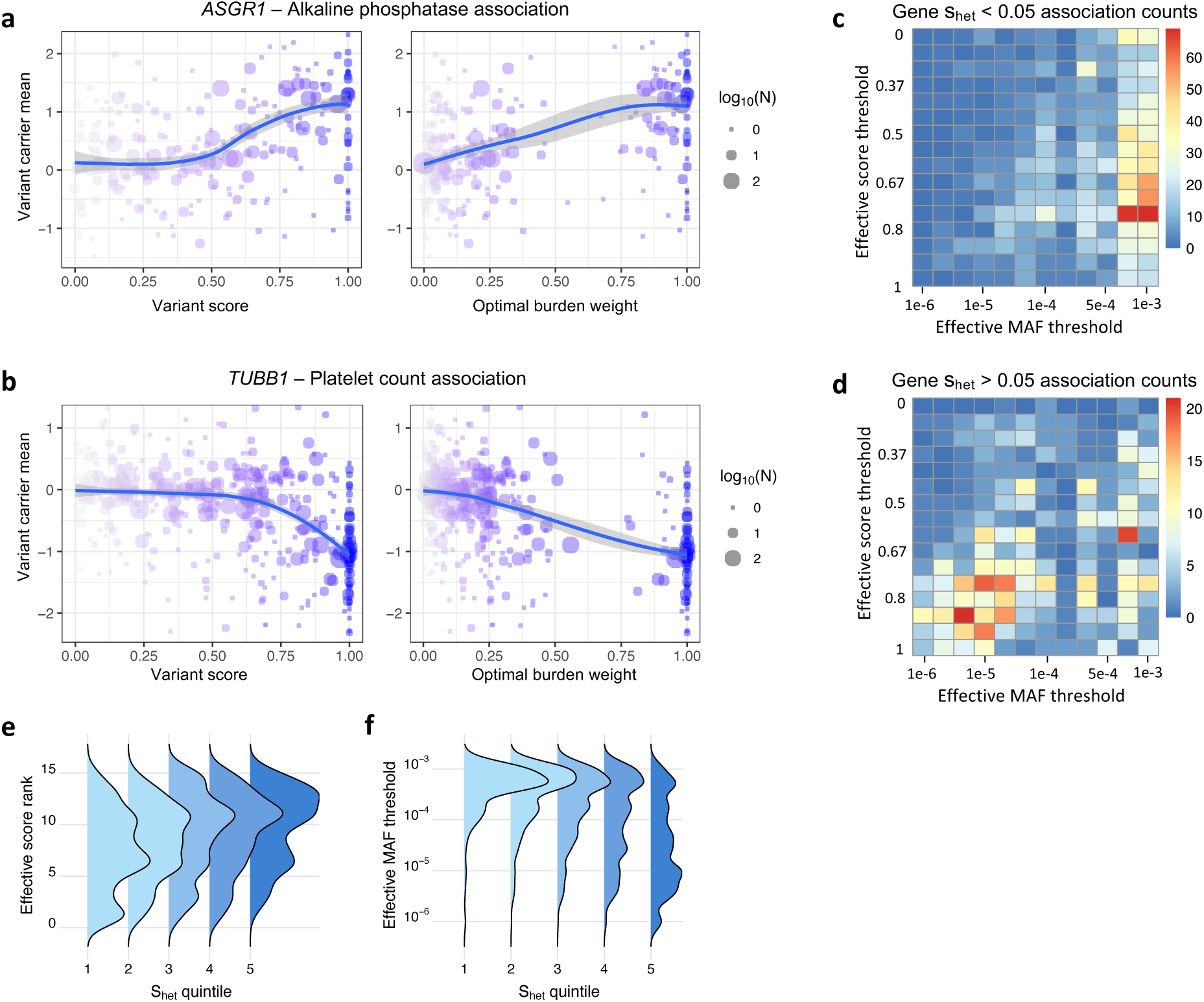
Nonlinear relationships between predicted variant pathogenicity and traits. **a**, Mean normalized alkaline phosphatase values for carriers of rare variants in *ASGR1* according to the variants’ predicted deleteriousness (PrimateAI-3D score) (left) or the variants’ weight from the most powerful burden test (right). **b**, Mean normalized platelet count values for carriers of rare variants in *TUBB1*. Blue curves show a loess fit to the data, with shaded 95% confidence intervals. **c,d,** Heatmap of the number of times each weight function had the smallest p-value among significant associations for genes with low constraint (**c**) or high constraint (**d**). **e,f**, Distribution of the effective thresholds on pathogenicity score (**e**) or MAF (**f**) for the optimal burden test weight function, with tested genes split into quintiles based on the s_het_ constraint metric.

Since different burden weighting schemes performed optimally in different genes, we investigated whether gene-level properties systematically influence the best-performing weights. To quantify this, we defined an “effective threshold” for each transformation, representing the cutoff at which a variant’s weight begins contributing significantly to the association test (**Methods**). For significant burden test associations, we visualized the number of times each weight function produced the smallest p-value. Stratifying genes based on estimated constraint or intolerance to damaging mutations in human populations (*s*_het35_) revealed that the best performing weight functions differ markedly between highly constrained and unconstrained genes (**Fig. 3c,d**). Specifically, as gene constraint increased, the optimal pathogenicity score threshold increased and the optimal MAF threshold decreased (**Fig. 3e,f**). A similar relationship was observed between increasing coding sequence (CDS) length and the optimal score threshold, while the relationship between CDS length and optimal MAF threshold was inconsistent (**Extended Data Fig. 10**). Although *s*_het_ and CDS length are correlated, in joint models both gene annotations remained significantly associated with optimal thresholds for score (*s*_het_ p = 1.1×10^-22^, CDS length p = 2×10^-30^) and MAF (*s*_het_ p = 1.2×10^-117^, CDS length p = 2.1×10^-6^).

When a gene is associated with multiple traits, a natural question is whether the individual rare variants have a similar impact across distinct traits. We found many examples where similar weights performed best across the trait associations of a given gene (**Extended Data Fig. 11-12**). Restricting to 46 quantitative traits with pairwise Pearson R^2^ < 0.3, we computed the intraclass correlation coefficient (ICC) of the effective pathogenicity score and MAF thresholds for each gene across traits. Among discoveries with FDR < 1%, pathogenicity score (ICC = 0.63) and MAF (ICC = 0.65) thresholds for each gene were highly consistent across their trait associations, suggesting that individual rare coding variants often have similar quantitative effects across traits.

### Nonlinear variant weights identify greater missense heritability and improve polygenic risk scoring

Previous work has suggested that LoF variants explain more heritability in rare variant tests than missense variants^30^. We investigate whether this proposition was consistent with FlexRV results, estimating the heritability recovered by burden tests with BHR across different weighting schemes and masks for 62 quantitative traits (**Methods**). To control for the possibility of overfitting on the same dataset, we selected FlexRV weights that produced p-values closest to but not lower than the omnibus CCT test p-value. Considering only missense variants, we found that the mean burden heritability recovered with FlexRV weights was 33% greater than that using a hard-threshold mask (PrimateAI-3D > 0.5, MAF < 0.1%), and 46% greater than using equal weights for all missense variants (**Fig. 4a**).

**Fig. 4:**
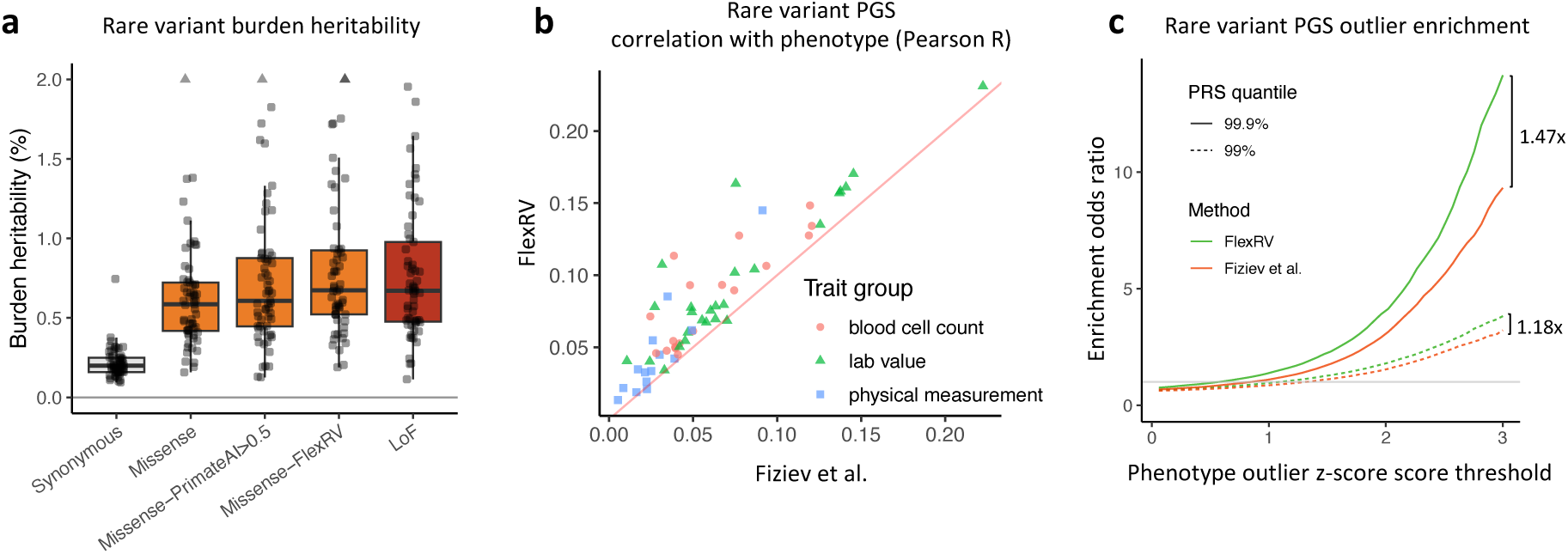
FlexRV applications to estimating burden heritability and developing polygenic scores. **a**, Burden heritability for rare missense variants is greater when using weights from FlexRV than when using a fixed cutoff or giving all missense variants equal weight and is comparable to LoF variant heritability. Each point is heritability within that variant category for a trait; triangles show heritability values truncated to 2% for plotting (2 traits). **b**, A PGS based on variant weights from burden testing in 256k UK Biobank individuals of continental European-ancestry demonstrates higher correlation (R^2^) with phenotype values in 64k independent continental European-ancestry UK Biobank individuals. **c**, Enrichment of individuals with high PGS scores (at 99% or 99.9% quantile) relative to all samples, among those with a phenotype value z-score above the given threshold.

Notably, the total burden heritability for rare missense variants was comparable to that for LoF variants when using FlexRV weights, indicating that using appropriate weights for missense and other types of variants will be key for exploring the genetic architecture of rare variants, especially when considering whole genome sequence data.

Lastly, we explored whether FlexRV-derived variant weights could improve polygenic scores (PGSs) for rare variants. Prior work^7,19^ has demonstrated that rare variant PGSs outperform common variant PGSs at identifying individuals with extreme phenotypes and exhibit greater cross-ancestry portability. To evaluate the impact of variant weights on PGS performance, we split 320,612 unrelated European ancestry individuals from UKB into 80% training and 20% test sets and performed rare variant burden tests in the training set using FlexRV weights or following the adaptive-threshold approach of Fiziev et al. We then modelled each quantitative trait as a simple linear function of the weight from the most powerful FlexRV burden test, or using PrimateAI-3D scores for variants included in the burden test from Fiziev et al. (**Methods**), and constructed rare variant PGSs from the linear model predictions. We found that rare variant PGSs derived from FlexRV weights consistently had equal or higher correlation (Pearson R^2^) with trait values in the held-out test set than PGSs from Fiziev et al. (**Fig. 4b**). Notably, individuals with high quantiles of FlexRV PGSs were also more highly enriched for having extreme phenotype values (**Fig. 4c**).

## Discussion

Studies of rare genetic variants have gained widespread interest due to their potential value in drug discovery, as targets with genetic evidence may succeed at a higher rate^36,37^, and allelic series of variants with different effect sizes may inform dose-response prediction^38^. Our method, FlexRV, increases the utility of genetic data by providing superior performance for gene discovery and phenotype prediction from rare variants. Although our approach allows for multiple, nonlinear relationships between variant annotations and complex human traits and disease, it is fully frequentist and unsupervised, allowing it to readily fit into most existing rare variant association testing pipelines. FlexRV is also feasible to run at biobank-scale: each FlexRV gene-trait omnibus test on ∼400,000 individuals took an average of 4 seconds for quantitative traits and 12 seconds for binary traits, requiring only 12-16 Gb of memory.

As the performance of pathogenicity predictors improves, it is possible that their scores will reflect phenotypic effects in a more linear manner, reducing the added benefit of our approach. There are reasons to believe this will not be the case. The best performing predictors to date leverage information encoded in the genomes of multiple species based on selection against protein-altering variants, as well as the variant’s protein structural context^17,33^. Nevertheless, natural selection acts on phenotypes, and established selection regimes such as stabilizing and disruptive selection have nonlinear effects^21^. More directly, the relationships between disease traits and their molecular and quantitative risk factors may be nonlinear due to developmental canalization^20^, gene-environment interactions^39^, or threshold effects^40^. Indeed, the relationships between diseases and risk factors, such as that between cardiovascular disease and either blood pressure or LDL cholesterol levels, may themselves be linear or nonlinear^41,42^. This implies that even if variant pathogenicity relates linearly to one trait, it may not to other traits, and considering nonlinear effects will likely remain important as pathogenicity predictors improve.

There are several directions for improvement of FlexRV. First, we have only explored burden and SKAT kernels. Considering other types of kernels, estimation procedures, or related nonlinear regression methods may further improve power^43–47^. Second, although we designed the set of 192 weights to reflect our biological intuition, alternative transformations may better approximate the true nonlinear relationships. Using additional, targeted weight transformations could result in additional discoveries, but the inclusion of additional tests reduces the significance of genes that do not benefit from these transformations. At some point, Bayesian approaches may provide a better tradeoff, but integrating these approaches^48,49^ into standard rare variant association test pipelines is more challenging than integrating FlexRV, which is both simple and computationally efficient. Moreover, given that optimal burden weights vary by gene constraint and gene (CDS) length, it may be possible to leverage gene properties when constructing weights to enhance power^35^. While we have not explored this in detail, we note that there is considerable heterogeneity even among highly constrained genes, where sometimes more lenient thresholds on predicted pathogenicity and MAF perform best. Finally, learning how to best integrate multiple variant annotations while allowing for nonlinear and variable effects across genes will likely further improve gene discovery.

**Extended Data Fig. 1:**
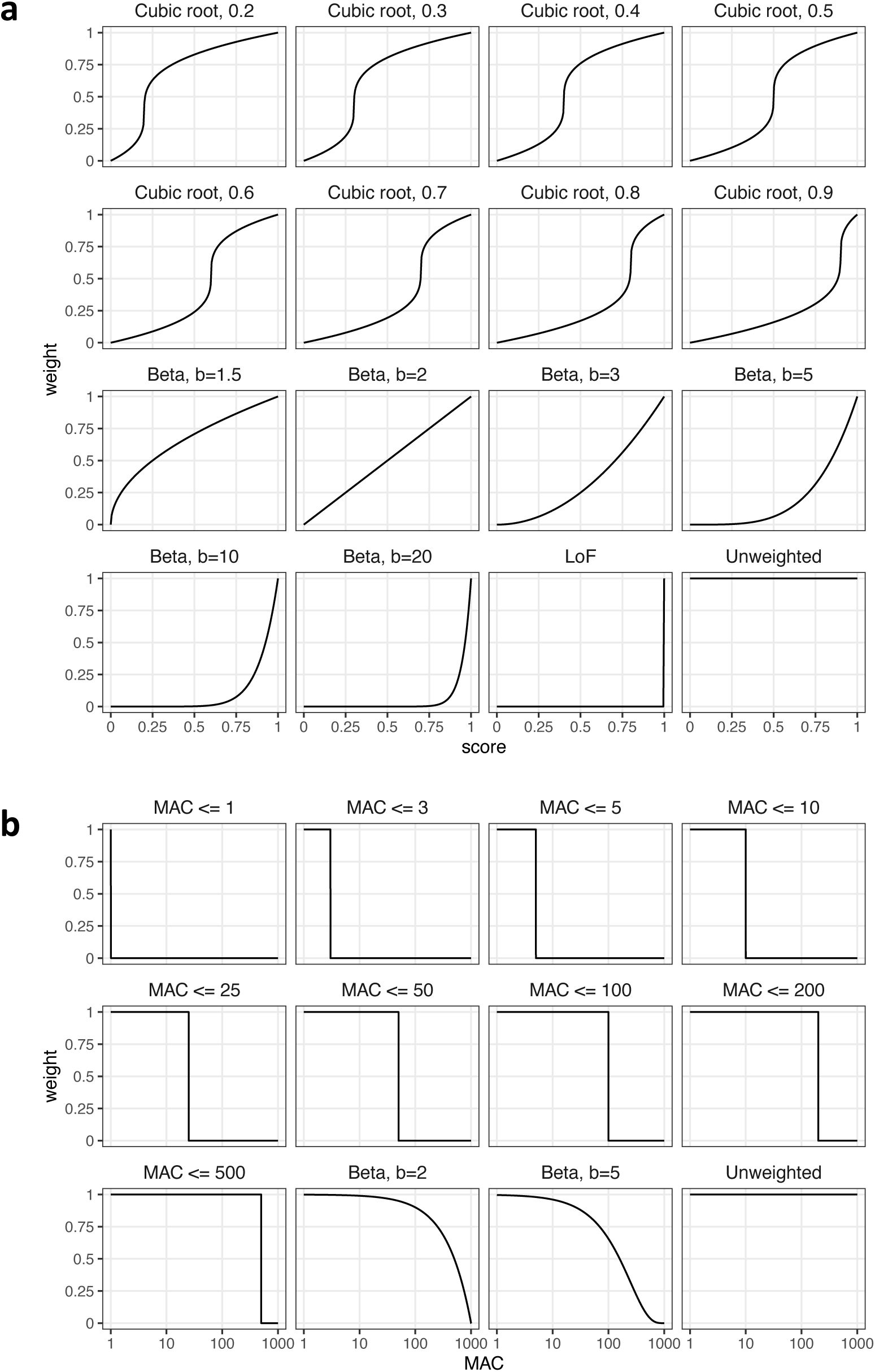
FlexRV weight functions. **a**, Transformations for the variant score (pathogenicity) annotation, which assume the score is defined on the interval [0, 1], and which output a weight in [0, 1]. **b**, Transformations for variant allele count, which assume an input allele count between 1 and 1000, and output a weight in [0, 1]. Beta distributions in both cases use parameters a = 1, and b as shown (Methods).

**Extended Data Fig. 2:**
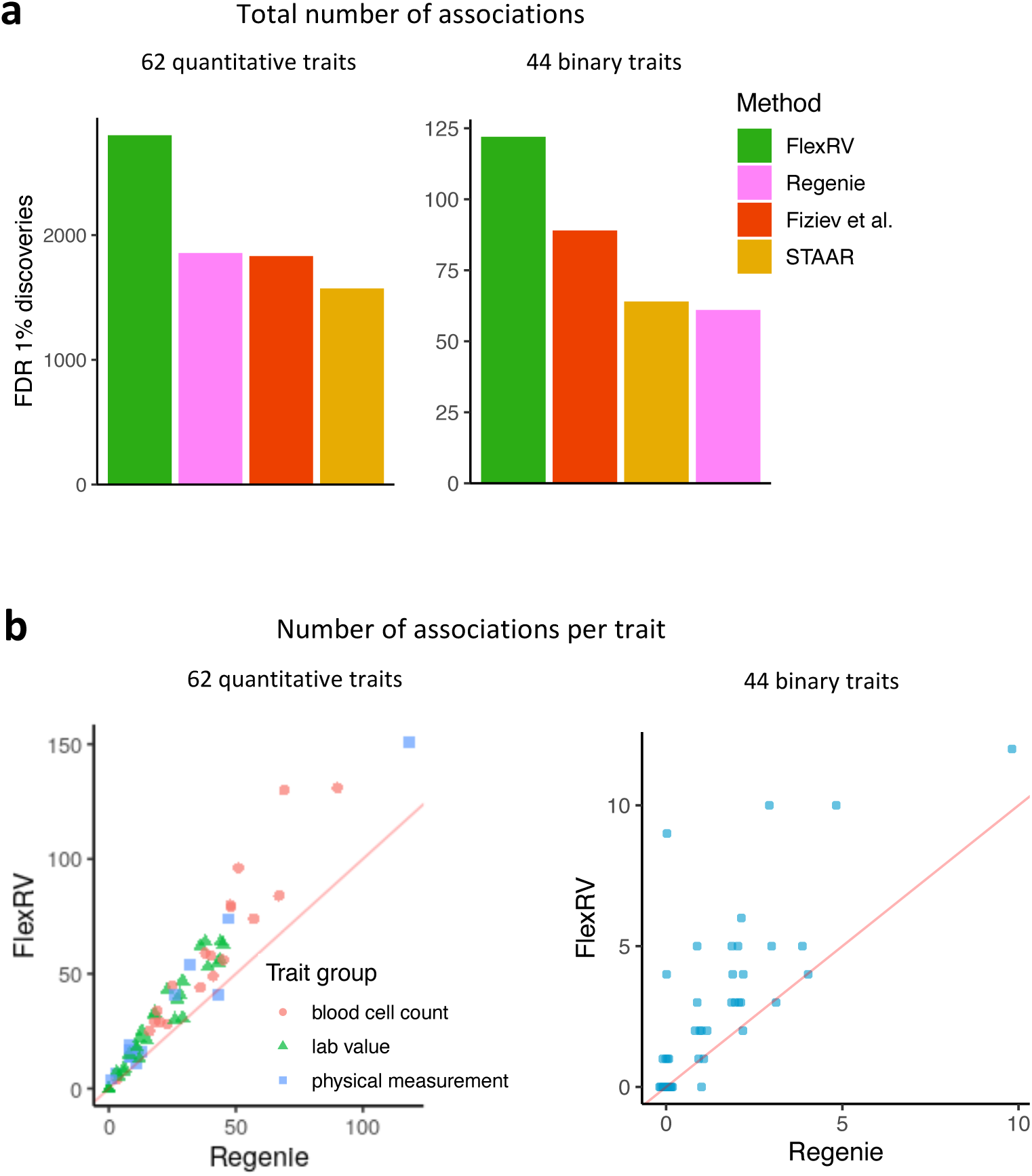
Number of associations for different burden testing methods. **a**, Total number of associations discovered at FDR < 1% for each method across 62 quantitative traits and 44 binary traits. **b**, Number of associations discovered at FDR < 1% for each trait, comparing Regenie and FlexRV, for 62 quantitative and 44 binary traits.

**Extended Data Fig. 3:**
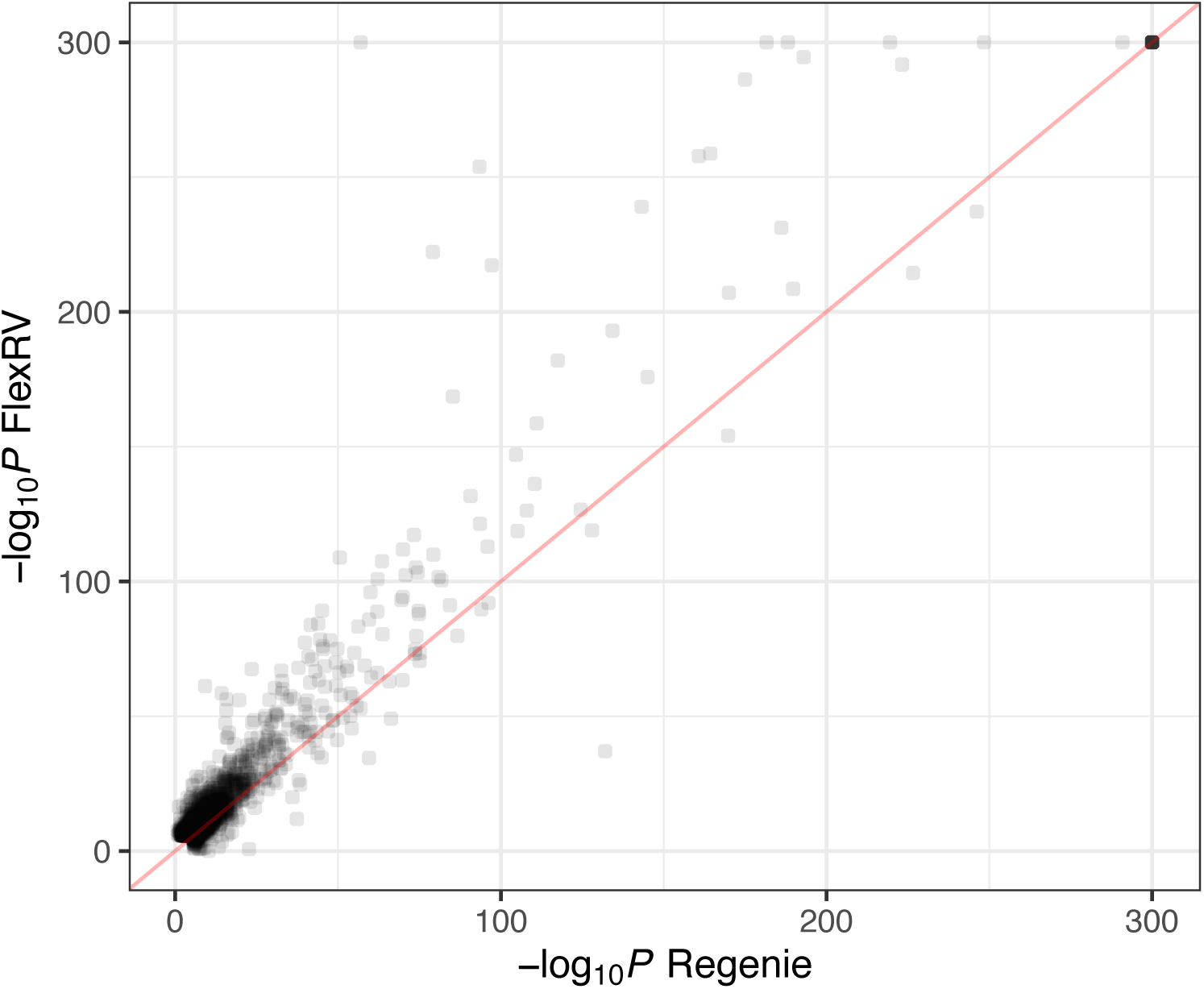
Scatterplot of -log_10_ p-values for FlexRV vs. Regenie. Each method finds some unique discoveries, but FlexRV shows more significant p-values for most genes called significant by either method.

**Extended Data Fig. 4:**
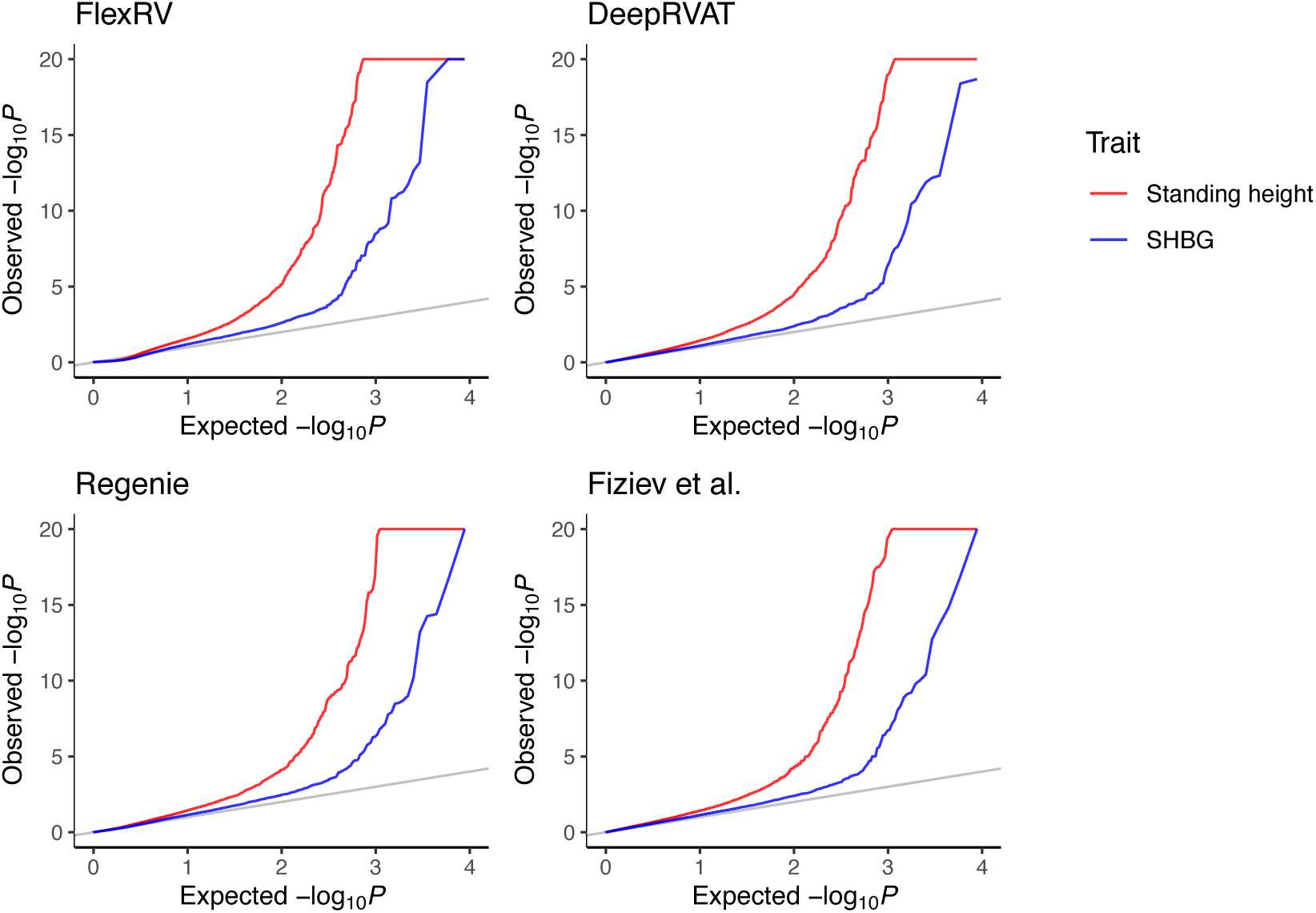
Quantile-quantile plots of FlexRV results for select highly powered traits. Shown are quantile-quantile (QQ) plots for the traits height and sex hormone binding globulin (SHBG), separately for each method. All methods show similar trends. Height is a highly polygenic trait, and the QQ plot shows an early rise above the “null expectation” of no association, suggesting that a large fraction of genes have some association. This is less true for SHBG, which appears less polygenic despite being well powered.

**Extended Data Fig. 5:**
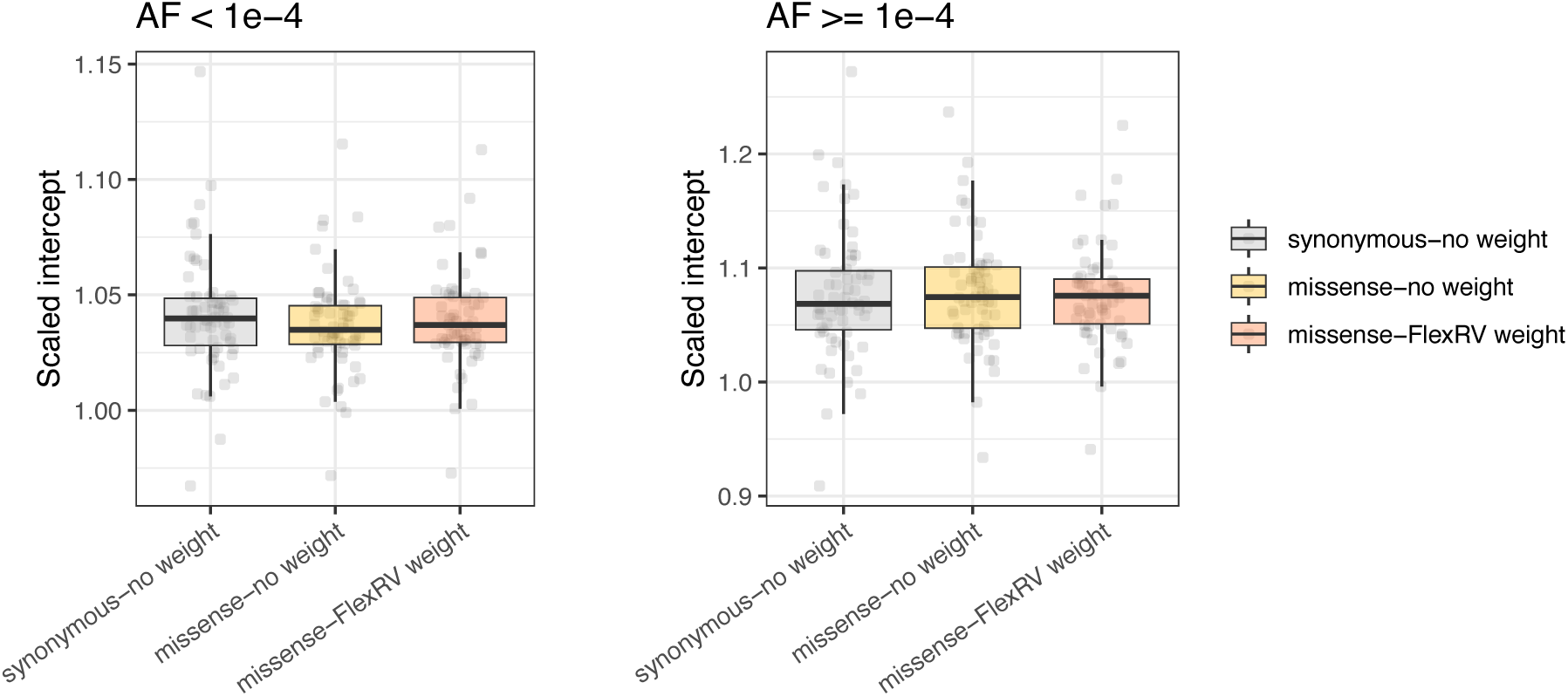
Burden heritability regression intercepts. Boxplots of the burden heritability regression (BHR) scaled intercepts for synonymous, missense (unweighted), or missense (FlexRV weights) across 62 quantitative traits, computed separately for ultrarare variants (MAF < 10^-4^) and for low frequency variants (10^-4^ <= MAF < 10^-3^). Intercepts are scaled as (intercept term / (1/n)) where n is the number of individuals. Estimates are within a similar range to that reported by in the original BHR publication on UK Biobank data^30^, providing evidence that inflation in the quantile-quantile plot above is primarily due to polygenicity rather than confounding.

**Extended Data Fig. 6:**
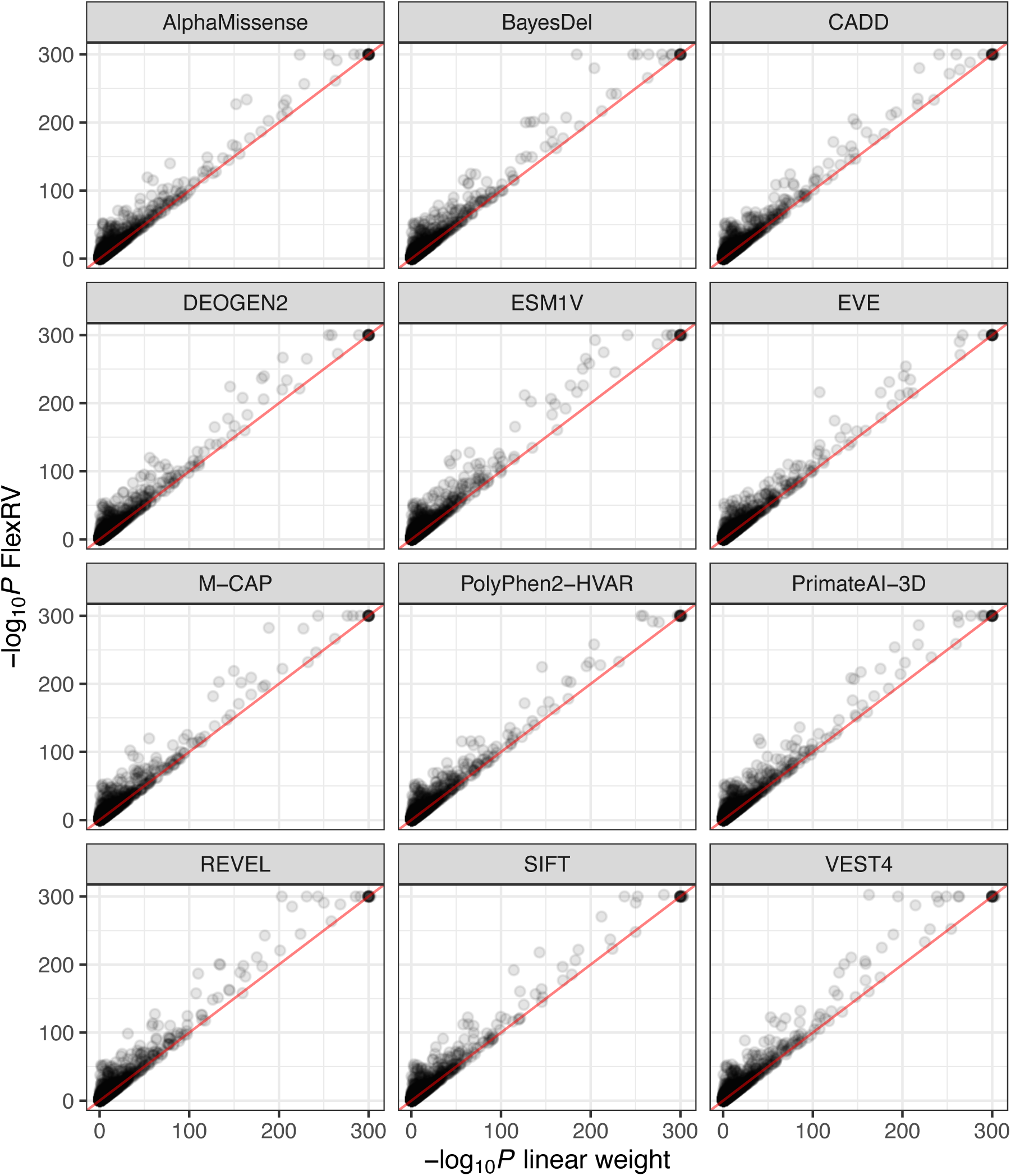
Improvement in p-values using FlexRV with different pathogenicity predictors. Each panel shows a scatterplot of -log_10_ p-values from burden tests using FlexRV (y-axis) vs. -log_10_ p-values from burden tests using the pathogenicity predictor as a (linear) variant weight (x-axis). In each case FlexRV tests were done with the indicated pathogenicity predictor and show results across 62 quantitative traits.

**Extended Data Fig. 7:**
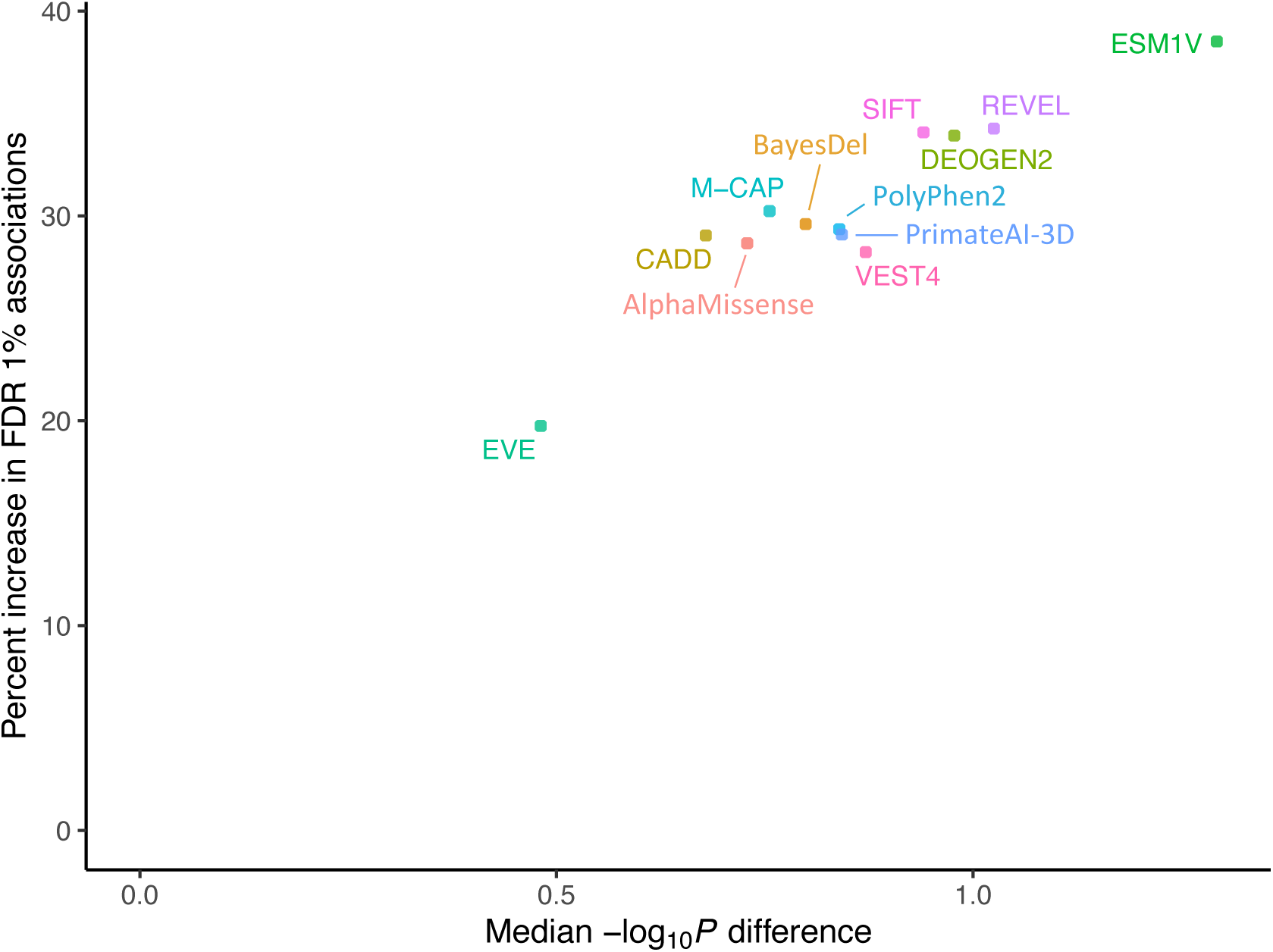
FlexRV yields improvements in association significance and number of discoveries. For each pathogenicity predictor we ran both FlexRV and burden tests using the predictor as a (linear) variant weight across 62 quantitative traits and considered gene-trait associations with FDR < 1% using either approach. The scatterplot shows the percentage increase in number of gene-trait associations using FlexRV relative to tests with a linear predictor weight (y-axis) vs. the median difference in -log_10_ p-values between FlexRV and the linear predictor weight. For all predictors FlexRV yields both more significant p-values and a greater number of associations.

**Extended Data Fig. 8:**
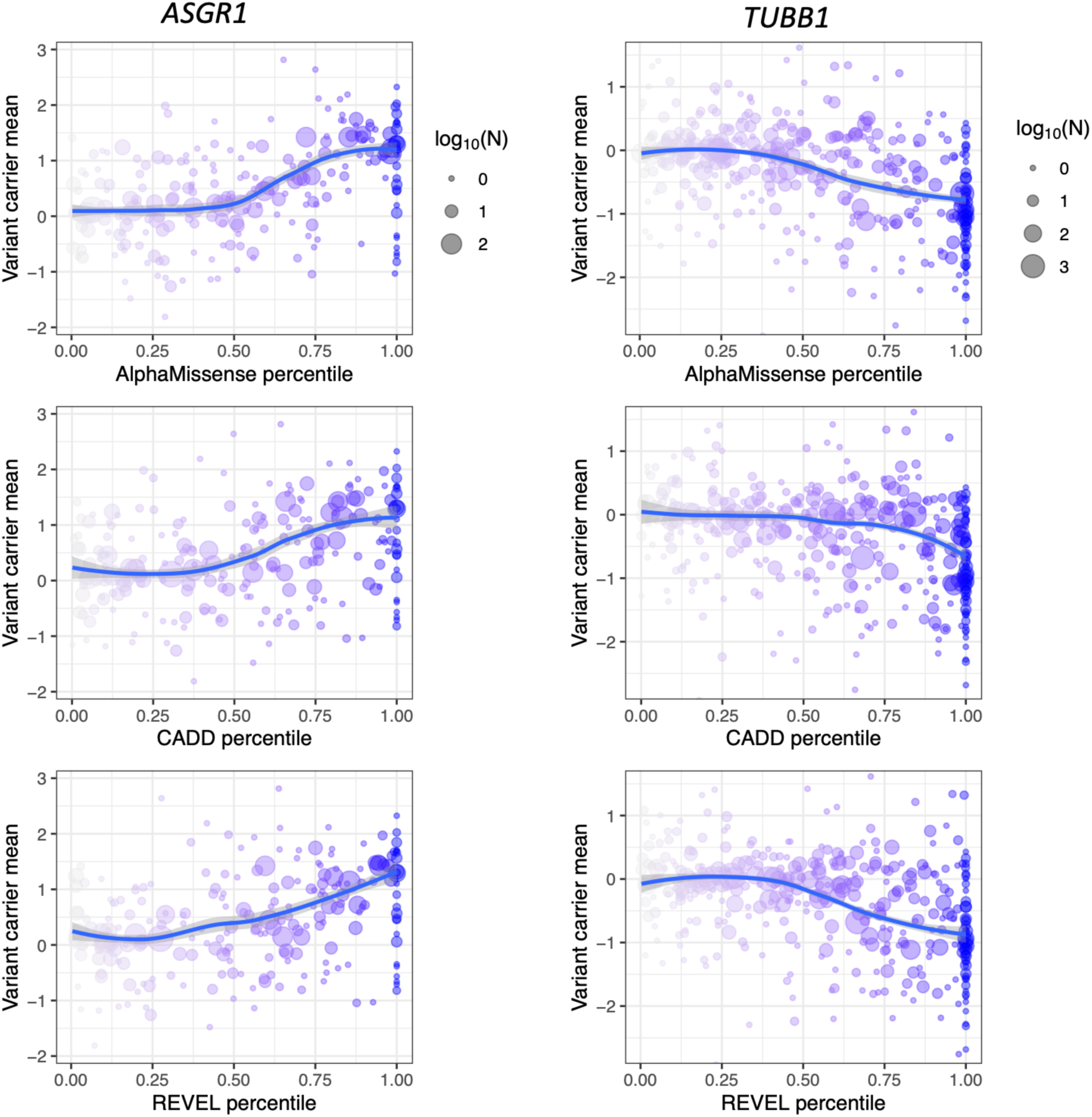
Variant pathogenicity scores have nonlinear relationships with phenotypes. Left panels show the mean normalized alkaline phosphatase values for carriers of rare variants in *ASGR1* according to the variants’ gene-based pathogenicity score percentile (from top to bottom: AlphaMissense, CADD, and REVEL). Right panels show the mean normalized platelet count values for carriers of rare variants in *TUBB1* according to the variants’ gene-based pathogenicity score percentile (from top to bottom: AlphaMissense, CADD, and REVEL). LoF variants are assigned a score of 1 in all panels. Point are colored by pathogenicity score (gradient from 0: grey to 1: blue). Blue curves show a loess fit to the data with shaded 95% confidence intervals.

**Extended Data Fig. 9:**
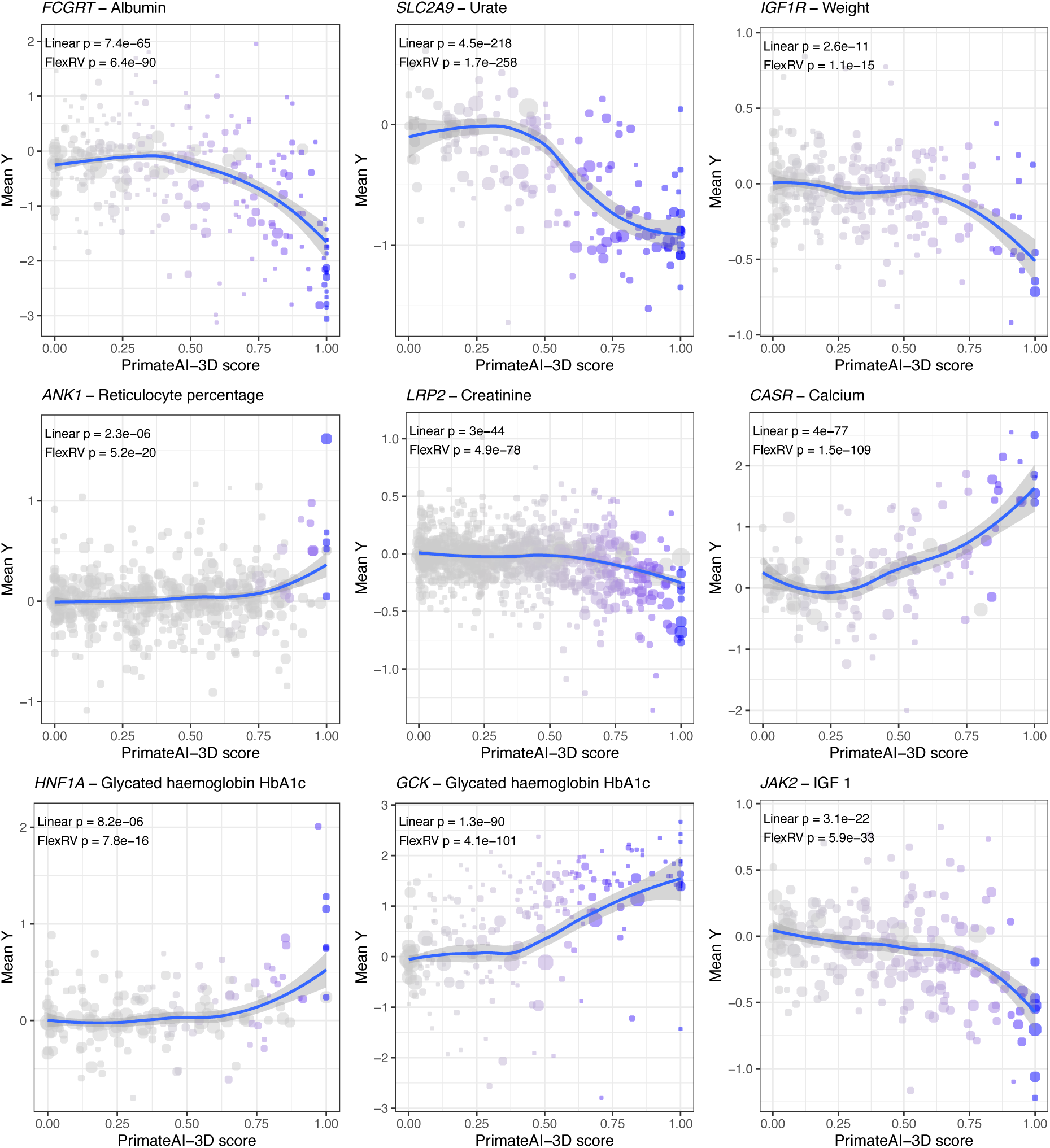
Examples of FlexRV nonlinear associations. Mean normalised phenotype values for carriers of rare variants for selected genes and traits compared to the variants’ PrimateAI-3D scores. Blue curves show a loess fit to the data with shaded 95% confidence intervals. Points represent variants, with size corresponding to log-scaled number of carriers, and blue color indicating the variant’s weight from the most powerful burden test.

**Extended Data Fig. 10:**
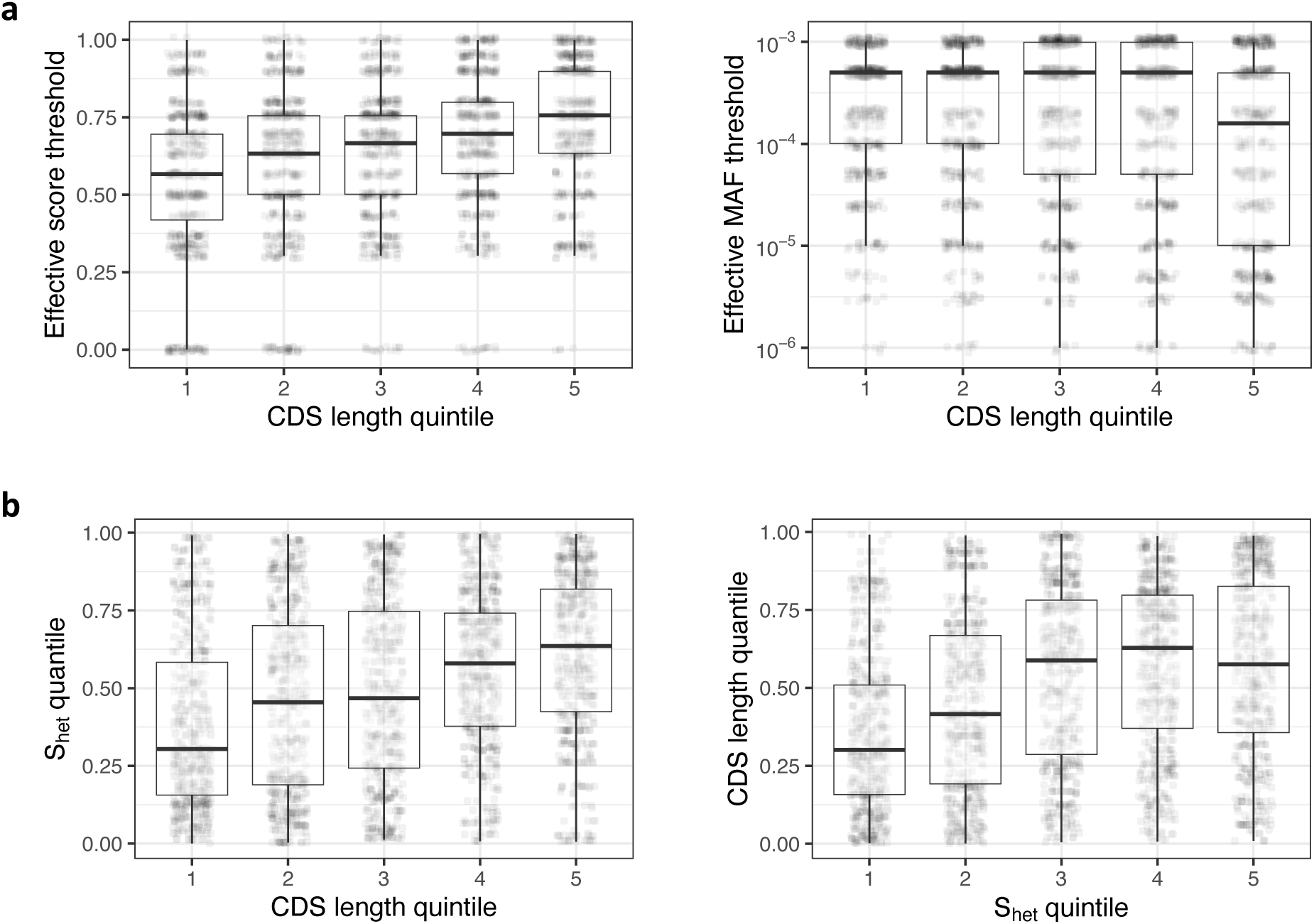
Correlations between gene CDS length, constraint (s_het_), and pathogenicity score thresholds among associated genes. All boxplots show values based on significant gene-trait associations. **a**, Boxplot of either the optimal score threshold (left) or MAF threshold (right), according to CDS length quintile of associated genes. The optimal score threshold increases with CDS length, but the optimal MAF threshold has an unclear relationship with CDS length. **b**, (left) Boxplot of gene s_het_ rank (quantile) according to CDS length quintile; (right) the same relationship but showing gene CDS length rank (quantile) according to s_het_ quintile.

**Extended Data Fig. 11:**
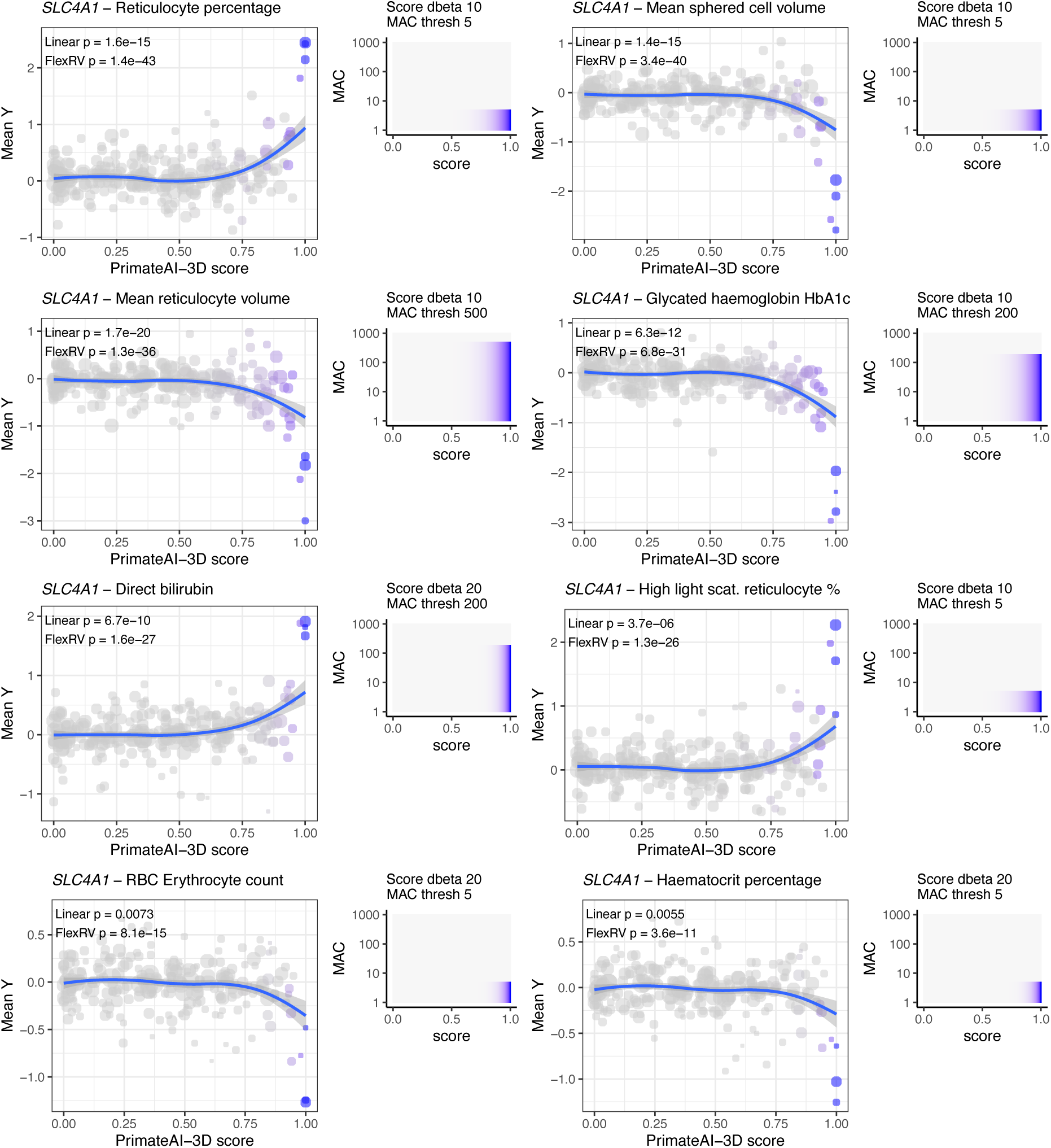
Distinct trait associations for *SLC4A1*. Panels show the top 8 trait associations for *SLC4A1*. Each panel shows: (left) a scatterplot of variant PrimateAI-3D score vs. mean normalised phenotype values for carriers of rare gene variants associated with the indicated phenotype, and (right) the weight function for the most powerful FlexRV burden test. Points represent variants, with size corresponding to log-scaled number of carriers, and blue color indicating the variant’s weight from the most powerful burden test. Blue curves show a loess fit to the data with shaded 95% confidence intervals. P-values in each panel indicate (top) the p-value from the single FlexRV test which used a linear weight according to variants’ pathogenicity score, with no weight on variant MAC (all MAC weights 1), or (below) the multiple-test-corrected p-value from FlexRV burden test across weight functions.

**Extended Data Fig. 12:**
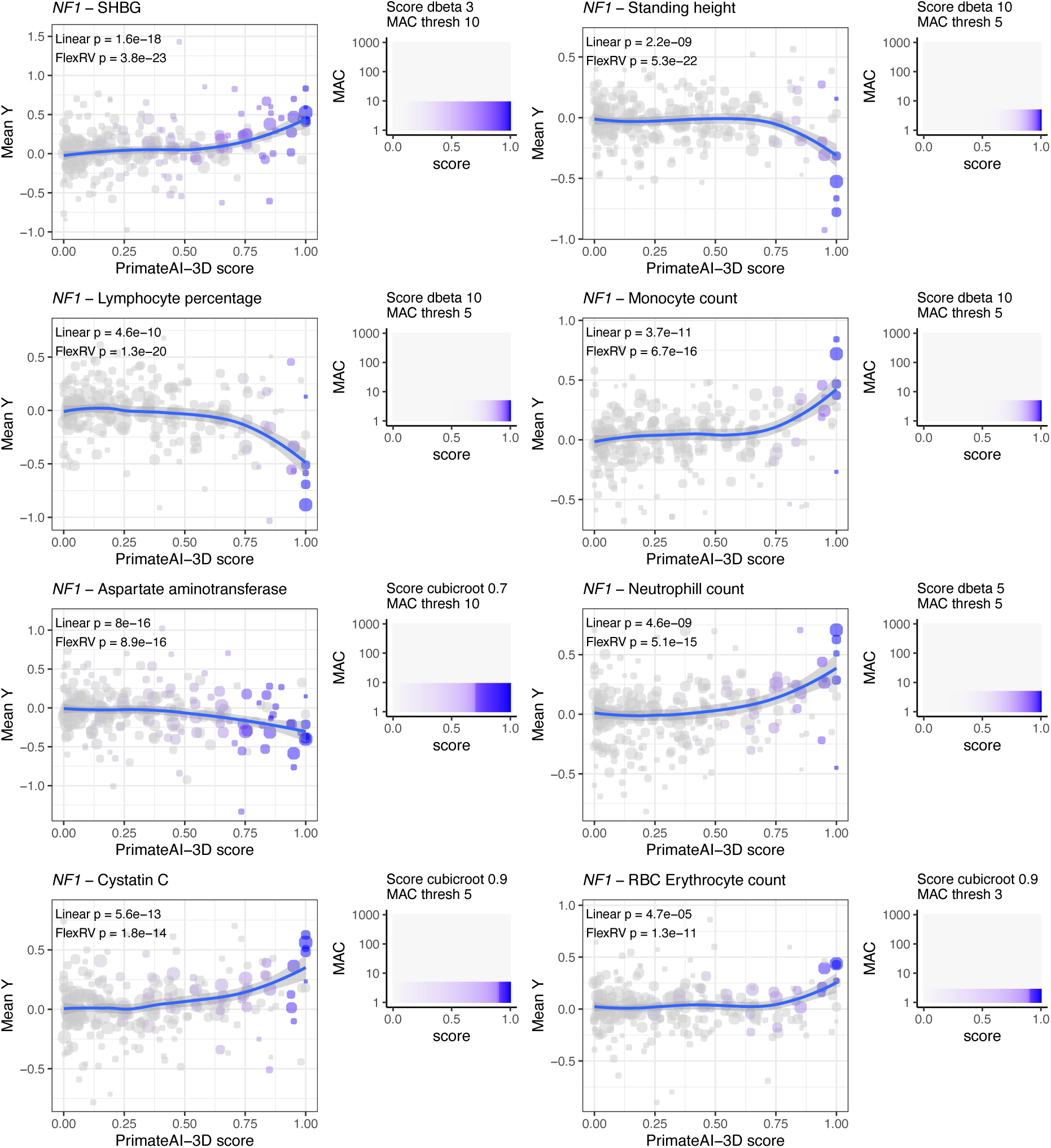
Distinct trait associations for *NF1*. Panels show the top 8 trait associations for *NF1*. Each panel shows: (left) a scatterplot of variant PrimateAI-3D score vs. mean normalised phenotype values for carriers of rare gene variants associated with the indicated phenotype, and (right) the weight function for the most powerful FlexRV burden test. Points represent variants, with size corresponding to log-scaled number of carriers, and blue color indicating the variant’s weight from the most powerful burden test. Blue curves show a loess fit to the data with shaded 95% confidence intervals. P-values in each panel indicate (top) the p-value from the single FlexRV test which used a linear weight according to variants’ pathogenicity score, with no weight on variant MAC (all MAC weights 1), or (below) the multiple-test-corrected p-value from FlexRV burden test across weight functions.

## Methods

### FlexRV variant weights

Consider the standard linear burden test model, E[**Y**|**G**,**X**] = **α** + **Gβ** + **Xγ**, where **Y** is a quantitative phenotype, **G** is an individual (n) x variant (m) matrix of rare variant carrier status (0, 1, or 2 alleles), and **X** is a covariate matrix. **α**, **β**, and **γ** specify the intercept, genetic, and covariate effects. Interest typically lies in evaluating evidence against the null hypothesis that no variants have any effect on the mean phenotype, H_0_: **β** = **0**. Typically, we set **β** = **1**β, forcing all variants to have the same effect under the null: this is how aggregation tests gain power over models fitting individual fixed effects to each variant. To allow for “individual” variant effects, we can *a priori* specify a relative weight for each variant, w_j_, which can be incorporated into the linear predictor by letting **Gβ** = **Gw**β, where each variant has an effect w_j_β on the conditional phenotype and interest is in testing evidence against the null hypothesis H_0_: β = 0. Notably, **w** is not a random variable in the standard burden test model and must be known ahead of time; using **Y**, **G**, or **X** to estimate **w** invalidates the use of asymptotic null distributions for common test statistics (*i.e.* burden and SKAT test statistics)^15,22^. We use this simple formulation to develop the intuition behind FlexRV.

If a burden test contains only LoF variants, it may be reasonable to assume that each variant has the same effect on the trait, and setting **β** = **1**β is a correctly specified model. However, other groups of variants such as missense and splice variants will likely have a distribution of effects. A recent model allows for estimating a single effect across groups of variants^50^, but we use the model above which allows directly specifying these relative effects for each variant as **w**. In practice, **w** is unknown and will almost certainly be misspecified, leading to a biased β̂ estimate. Most importantly for gene discovery studies, our test will be underpowered when β ≠ 0. Previous approaches have approximated **w** based on MAF, genomic annotations, and pathogenicity predictions^7,14,18,22^. Some recent methods use multiple predictors or annotations to obtain a set of **w** vectors, run a burden test with each, and combine the results, controlling for the multiple tests performed^14,18^. Notably, the principle behind this type of approach is that the power gained by (more) closely specifying the correct aggregation test model at some number of genes is greater than the power lost after adjusting for the multiple tests performed.

The core idea of FlexRV is that a variant annotation or pathogenicity prediction may require a nonlinear transformation to better match the true effects of a set of rare genetic variants on a complex trait of interest. In general, we have no expectation that a particular annotation or predictor *should* be linear in all complex phenotypes of interest, but we may often hope for a monotonic or smooth relationship. Since learning this **w** with in-sample data invalidates the assumptions of broadly used generalized linear mixed model (GLMM) and kernel regression implementations^51^, we choose instead to apply a set of prespecified transformations to an annotation or predictor, **s**. We use each transformed weight **w** = *f*(**s**) as the relative effects of variants in our rare variant test, and combine the results of all tests.

We start by combining a single annotation each for LoF, splice, and missense variants into a single score. For putative loss-of-function variants (frameshift insertions or deletions, stop gain, stop loss, start loss, canonical splice variants), we assign the highest score of 1, since these are likely the most pathogenic. For missense variants, we opt to use the pathogenicity predictions from PrimateAI-3D, which we have previously shown to have high performance in distinguishing between pathogenic and benign rare variants^17^. We obtain the raw PrimateAI-3D score for each observed variant and convert this to a percentile across all possible (but unobserved) missense variants in its gene. To create our initial score, **s**, we combine the vectors for LoF and missense variants, taking the max in the rare case of multiple scores.

To approximate the relationship between this score and a complex phenotype of interest, we applied 16 transformations, *f*(**s**), to this score as follows (see **Extended Data Fig. 1**):

- Unweighted (**w** = **1**)
- LoF only (w_j_ = 1 if LoF, otherwise w_j_ = 0)
- Cubic root with each of 8 different transition values
- Beta distribution functions with each of 6 different shape parameters

Cubic root functions, *f*, were defined to map input scores, **s**, on the interval [0,1] to weights, **w**, in [0,1], with a transition based on a “transition” value, t, selected from {0.2, 0.3, 0.4, 0.5, 0.6, 0.7, 0.8, 0.9} as follows:

**w** = (sign(**s**−t)⋅|**s**−t|^1/3^ + t^1/3^) / (|1−t|^1/3^ + t^1/3^)

Beta distribution functions, *f*, also mapped input scores, **s**, on the interval [0,1] to weights, **w**, in [0,1], based on the beta probability density function, dbeta, with fixed parameter a = 1, and parameter b selected from {1.5, 2, 3, 5, 10, 20} as follows:

**w** = dbeta(1 - |**s**|, 1, b) / dbeta(0, 1, b)

We also applied 12 transformations to variant minor allele frequency (MAF):

- Unweighted (**w** = **1**)
- Beta distribution functions with 2 different shape parameters
- Hard threshold functions such that w_j_ = 1 for variants at or below the threshold and w_j_ = 0 otherwise

Beta distribution functions, *f*, followed the definition above, but we mapped input scores **s** to **w** as *f*(**MAF** / 0.001). This scales **w** such that the most common variants included in this test will have the smallest scores (∼0). Parameter b was selected from {2, 5}.

Threshold functions, *f*, map input scores, **s**, to weights **w** based on MAF thresholds, where the MAF thresholds were chosen to correspond to minor allele counts (MAC) in {1, 3, 5, 10, 25, 50, 100, 200, 500}.

After obtaining a weight vector from each transformation based on the pathogenicity score or annotation, **w_s_**, and the transformation based on MAF, **w_MAF_**, we obtained a final **w** = **w_s_w_MAF_** by taking the product of the two scores, resulting in a set of 192 weight vectors.

We perform one rare variant test (or more if additional kernels are used, e.g. SKAT) for each gene-trait combination for each weight vector, and obtain a combined p-value, p_FlexRV_, that controls for multiple testing (at least in the tail) by applying the Cauchy Combination Test (CCT) to the set of p-values, p_1_, …, p_k_, as follows:

Construct the test statistic, 𝑇 = ∑_*i*_ tan(0.5 − 𝑝_*i*_) 𝜋

Compute the p-value, 𝑝_*FlexRV*_ = 𝑃[𝐶(0,1) ≥ 𝑇], where 𝐶(0,1) is a Cauchy distribution

### FlexRV implementation

We implemented FlexRV in R (v4.3.3) using the STAAR package^14^ to perform statistical tests, which fits generalised linear and mixed models (GLMMs) for quantitative or dichotomous traits, allowing for sparse genetic relationship matrices (GRMs) that account for relatedness and population structure. In this study, we used unrelated individuals and so we did not make use of GRMs, but it is straightforward to do so with FlexRV. We reimplemented the STAAR function to apply the weight transformations as defined above, rather than those originally defined by the authors. We called STAAR with our sets of weights and combined the results of the tests using CCT^10^, which controls for multiple testing. The authors of DeepRVAT^19^ reported that STAAR is miscalibrated for binary traits with imbalanced case-control ratios; subsequently the STAAR authors added an implementation of the saddlepoint approximation (SPA) (v0.9.7), which provides substantially less biased estimates in this scenario. Therefore, for binary traits we also ran STAAR, with SPA max_iter parameter of 20, which we found to give well calibrated results in permutation tests.

We found that STAAR ran slowly for genes with a large number of variants. Therefore, for genes with >= 400 variants we collapsed variants having total AC <= 5 into up to 50 pseudo-variants based on each AC from 1 to 5, and pathogenicity score in bins of 0.1 (i.e. 0 - 0.1, 0.1 - 0.2, …, 0.9 - 1.0). Each pseudo-variant represents multiple variants with the same AC and was assigned the mean pathogenicity score of variants in the bin. The association p-values obtained from burden tests with binning were highly similar to those without binning. We note that the idea of pseudo-variants has been used before in the context of rare variant testing^13^, but our approach allows for more flexible grouping and assignment of scores to pseudo-variants which likely results in increased power. More generally, we note that a binning approach such as the one implemented here may protect against the effects of model misspecification^13^.

### UK Biobank data

We used genotype data from whole exome sequencing of 454,712 individuals released by the UK Biobank in October 2021, as described in Fiziev et al^7^. We excluded individuals who had more than second degree relatedness by removing one individual of each pair who had a UK Biobank-provided kinship coefficient greater or equal to 0.0884. To maximize power in our burden tests, we kept individuals from all ancestries, which gave us 423,614 unrelated individuals. We used common and rare variant principal components as computed by Fiziev et al.

We selected a broad range of 62 well-powered quantitative phenotypes, including anthropometric traits, blood cell counts, and blood and urine biomarkers (**Supplementary Table 1**). For systolic blood pressure we took the average of the two consecutive measurements at each visit. For spherical equivalent, we first computed the values for the left and the right eyes as “spherical power” + 0.5 × “cylindrical power” and then took the average between the two eyes as the phenotype value for each person. We inverse rank normal transformed (IRNT) phenotype values for use in association tests.

Disease phenotypes were derived from category “First occurrences – Health-related outcomes” (category 1712) and were set to 1 if a value was present in the field “date XX first reported” (where XX is the ICD10 code for the disease), and 0 otherwise.

### Variant annotation

We used the variant effect predictor (VEP) v109.3 to annotate variant consequences^52^. For variants with multiple consequences, we kept the single most severe consequence. We excluded variants for which more than 5% of individuals had missing genotype calls, and we excluded in-frame insertions and deletions due to uncertainty in how to assign a deleteriousness score. We annotated missense variants with PrimateAI-3D^7^ and 11 additional pathogenicity predictors obtained from dbNSFP^53^, and we considered both raw scores and scores corresponding to the percentile of the variant’s score among all possible missense variants in the gene. Initial testing showed that transformed scores gave equal or greater power in burden tests with the added benefit that score distributions were consistent across genes and predictors. While this may change the shape of the relationship between pathogenicity scores and phenotypes (which are often transformed themselves, *i.e.* IRNT), we note that we frequently observed nonlinear relationships with both the original and transformed scores. We therefore used gene-based pathogenicity score percentiles in all subsequent analyses. After annotation, we retained variants in 18,790 unique protein-coding genes (based on HGNC symbol) with MAF < 0.1% in the full cohort for gene burden tests.

### FlexRV burden tests

For tests with quantitative traits, we used a two-stage procedure to account for covariate effects^54^. First, we standardized phenotypes using IRNT. We selected covariates: age, sex, age*sex, UK Biobank assessment center, 20 common variant genotype PCs, and 20 rare variant genotype PCs, as well as a common variant polygenic score (PGS) as a covariate. We defined this PGS based on GWAS for the trait, for which we had identified independent signals in on Fiziev et al^3^. Briefly, we used ridge regression with 5-fold cross-validation to fit phenotype values based on these associated common variants, and used the model fit to derive a PGS for each individual. Using STAAR, we fit a null model with all covariates, obtained residuals from this model, and re-transformed these values using IRNT. We used these IRNT-transformed residuals as the phenotype of interest and re-fit the null model that again included the above covariates. To compute the score test, we compared a “full” model with covariates and sample genotypes to this “null” model with only covariates. For each gene tested and each FlexRV weight function, we determined the number of individuals with a nonzero weight variant, and if there were fewer than 5 individuals we excluded the weight function.

For tests with binary traits, we used case (1) and control (0) values as defined above as the phenotype for logistic regression with the same covariates as for quantitative traits, again computing a trait-specific common variant PGS. We first tested each gene giving STAAR parameters for SPA of max_iter = 10 and p_filter_cutoff = 0.01. If the gene’s p-value was less than the cutoff after 10 iterations, we re-ran the test with max_iter = 200. This improved computational efficiency since for most genes 10 iterations was sufficient for accurate p-value estimation, but it ensured that more iterations were used if needed.

### Permutation and synonymous burden tests

To run FlexRV with permuted data for quantitative traits, we first fit covariates and IRNT-transformed the residuals as described above. Assuming the null model is specified correctly, these values are exchangeable (under the null), and we swapped phenotype values at random to obtain permuted values. In total, we permutated phenotype values 5 times for all genes for 62 phenotypes and ran FlexRV. To run FlexRV with synonymous variants, we need to assign a score to each variant. Although the simplest approach is to equally weight each variant and assign a score of 1 to each variant, we wanted to better mimic the FlexRV procedure and assigned scores by randomly sampling, with replacement, from the PrimateAI-3D scores for missense variants in the same gene.

### STAAR burden tests

We ran STAAR^14^ using the same variant and phenotype data as for FlexRV, and the same null model including covariates. We re-implemented the STAAR and STAAR_Binary_SPA functions with a fix so that any “NA” p-values from a set of annotation weights were ignored before combining p-values with CCT, which rescued some genes that would otherwise have no result. We used the FAVOR database^26^ to annotate variants with annotation PCs as described in the study introducing the STAAR method^14^. Specifically, from FAVOR we used apc_conservation_v2, apc_epigenetics_active, apc_local_nucleotide_diversity_v3, apc_mappability, apc_micro_rna, apc_mutation_density, apc_protein_function_v3, apc_proximity_to_coding_v2, apc_proximity_to_tsstes, apc_transcription_factor. For each annotation PC, as required for input to STAAR, we converted to a Phred-scaled score as −10 × log_10_(rank(−score)/M), where M is the total number of UKB exome variants.

### Regenie

Using plink2^55^, we filtered array-based genotype data from UK Biobank to MAF > 1%, pruned variants using a window of 200kb and R^2^ threshold of 0.8, and thinned them to 500,000 total variants. We used these data to run the whole-genome regression Regenie^23^ step 1 for 62 quantitative traits, and separately for 44 binary traits, with a block size of 1000 and 5-fold cross-validation. We annotated the same rare variants (MAF < 0.1%) used in FlexRV tests with the 5 pathogenicity predictors used in Backman et al.^9^, SIFT4G^56^, Polyphen2 HDIV and HVAR^57^, LRT^58^, and MutationTaster^59^, obtained using dbNSFP^53^. For simplicity, we limited tests to genes on autosomes (*i.e.* excluded chrX). For burden tests (Regenie step 2), we defined 8 variant masks consisting of 4 MAF thresholds (0.1%, 0.01%, 0.001%, singleton) applied to either LoF variants only, or LoF variants and those for which at least one of the predictors scored the variant as deleterious. We also tried masks that included missense variants called deleterious by all 5 predictors, as in Backman et al.^9^, but this resulted in much lower power, and so we used the more powerful mask for Regenie in our comparisons. We used the same covariates as for FlexRV in all steps with Regenie, except that we did not include a common variant PGS. For binary traits we used SPA correction with a p-value threshold of 0.01. We combined the different Regenie mask results using CCT to obtain a single p-value per gene-trait pair for comparison with other methods.

### DeepRVAT

We downloaded precomputed DeepRVAT^19^ association data from https://doi.org/10.5281/zenodo.12736824, filtering to results from the “470k_aa” cohort. For comparisons with DeepRVAT, we included all 28 quantitative traits tested by both methods. For binary traits, DeepRVAT used trait definitions that sometimes included multiple ICD codes, in contrast to our analyses. We therefore used the 24 binary traits that were likely to be an exact match due to being defined by a single ICD code for comparison (**Supplementary Table 1**). When comparing association testing methods, we used Benjamini-Hochberg FDR adjustment separately for each method and trait to identify associations significant at FDR 1%.

### Method comparisons

In all comparisons of methods, including power, replication, and enrichments, we subsetted results to autosomal gene-trait pairs where a successful test was available for all methods. For comparisons involving Regenie, STAAR, and Fiziev et al. this removed a modest fraction (0.25%) of gene-trait pairs where at least one method did not compute a result. For comparisons that also included DeepRVAT, 0.6% of gene-trait pairs were removed (or could not be matched).

### Pathogenicity predictor comparisons

To assess FlexRV performance with different pathogenicity predictors (**Extended Data Fig. 6-8**), we annotated missense variants with dbNSFP^53^ and ran FlexRV on variants with MAF < 0.1% using weights from each predictor for 62 quantitative traits in the UKB exome cohort. We compared FlexRV burden test results based on all weight transformations (multiple test-corrected) to those from burden tests using the pathogenicity prediction as a linear weight with no additional MAF cutoffs.

### Replication – downsampling

To compare results from a smaller sample set to the full cohort, we first randomly sampled 200,000 individuals from the unrelated individuals in our main analysis, and ran burden testing on these individuals using FlexRV, STAAR, and Regenie. For Regenie, we downsampled the array genotypes to these individuals for Regenie step 1, and downsampled the exome data for step 2 burden tests. To evaluate methods, we used DeepRVAT results from the full 470k cohort as a truth set, since the annotations and approach are largely independent of the other methods. We assigned as true positives genes called significant by DeepRVAT at a Bonferroni threshold (p < 0.05 / N) per trait, where N is the number of genes tested per trait, and all other genes were considered negatives. We ordered genes by p-value separately for each method and plotted the cumulative number of replicated genes as maximum gene rank increased (**Fig. 2a**).

### Replication – All of Us

The All of Us (AoU) research program^31^ is a longitudinal cohort supported by the National Institute of Health aiming to recruit a diverse group of at least 1 million individuals across the USA for biomedical and health research. Our use of AoU data was approved under a data use agreement between Illumina and the AoU research program. For evaluating replication of our burden test discoveries made in UKB, we used the gene-based cross-population meta-analysis rare variant results of the “All by All” v1 tables provided by AoU (downloaded Sept 23, 2024), which were computed using standard burden tests based on WGS of up to ∼250,000 individuals. For each of our 62 quantitative UKB phenotypes, we manually searched the AoU data browser and identified the OMOP concept ID that matched best, if available. We then used the 28 available matching phenotypes which had at least 5,000 samples in AoU and which had at least one gene burden test with p < 1×10^-6^. We used CCT to combine p-values for each gene across the 9 distinct pLoF and missense masks that were tested (ignoring the synonymous and “Cauchy” masks). Due to the relatively lower power for association detection in All of Us compared to UKB, we defined replication of a UKB burden test association as those having All of Us p_CCT < 0.05 for the same gene-trait pair.

### GWAS nearby genes and PoPS enrichment

We used GWAS data computed by Fiziev et al.^7^ on the same UKB samples and traits, for which independent signals at genome-wide significance (p < 5×10^-8^) had been identified. We identified all genes with start or end coordinates within 500kb of each GWAS index variant. We assigned locus-based ranks to genes based on their minimum distance to the index variant and limited the table to the nearest 20 genes per index variant. To determine the GWAS enrichment for different burden testing methods, we ordered burden test results for each method by p-value across all 28 traits available for all methods, and selected the top 2,500 gene-trait associations for each method. By using a fixed number of associations, we avoid biasing enrichment results based on differences in where the threshold of “significance” is determined by different methods. We used different definitions of “GWAS hit gene” based on whether the gene is among the nearest 1, 2, 5, or 20 genes to the GWAS index variant, referred to as G_k_ below, k ∈ {1,2,5,20}. We then constructed 2×2 tables contrasting the number of GWAS hit genes among the top 2,500 burden associations to the number in the bottom 50% of burden associations across all traits, and determined the odds ratio (OR) for top 2,500 genes to be GWAS hits using Fisher’s exact test. That is, for A = sum(G_k_) in top 2500, C = sum(G_k_) in bottom 50%, and with N = number of bottom 50% genes, the OR is computed as (A)(N-C)/[(2500 - A)(C)].

For each trait we ran PoPS^3^ v0.2 with the original features used by the authors (https://github.com/FinucaneLab/gene_features), but excluding KEGG pathways. We defined PoPS hit genes to be those in either the top 0.5%, 1%, 2%, or 5% of PoPS scores genome-wide for each trait, and similarly determined enrichment of PoPS hit genes in the top 2,500 burden associations relative to the bottom 50% analogous to the enrichment for GWAS.

### Relationship of weight functions to gene properties

Our score transformations assign a different weight to each variant based on its pathogenicity score. To define an “effective threshold” for each transformation function *f*, we computed the integral of 1-*f*:

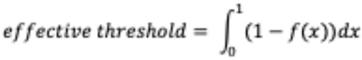

This definition corresponds to the mean unweighted variant score for each *f* – in other words, variants with scores below this value receive relatively little weight (after transformation). For example, the effective score threshold for a cubic_root function with shift 0.7 (value at which f(x) = 0.5) is equal to 0.7. We defined effective MAF threshold in the same way for beta functions and assigned the actual MAF cutoff for threshold functions. We counted the number of times each weight function resulted in the lowest p-value across all quantitative trait tests and visualized this as a heatmap ordering the weight functions by effective score or MAF threshold in **Fig. 3**. To relate effective thresholds with gene properties, we obtained s_het_ scores from Supplementary Table 1 of the GeneBayes^35^ study. We also obtained gene constraint values (and CDS length) from gnomad v2.1.1^60^ (https://gnomad.broadinstitute.org/data#v4-constraint), and found highly similar results to those using s_het_. To determine whether both are independently related to the best-performing FlexRV transformation thresholds, we used linear models predicting the score (or MAF) threshold value from gene s_het_ rank and CDS length rank.

To compare weights across traits, we first obtained an approximately independent set of phenotypes by computed the pairwise correlation of all 62 quantitative phenotypes, and then removed one of each pair for which R^2^ > 0.3, generally keeping the trait which had more burden associations. Using this set of 46 traits, we then computed the intraclass correlation (ICC) of effective score and MAF thresholds for each gene across traits using the performance^61^ and lme4^62^ R packages, with R v4.3.3.

### Burden heritability

To estimate the trait heritability captured by burden tests, we used burden heritability regression (BHR)^30^. As input for BHR, we computed single-variant association statistics for MAF < 0.1% variants, using 2-sided *t*-tests comparing rare variant carrier phenotypes to those of non-carriers, across all 62 quantitative traits. Phenotype values were the residuals from null models including the same covariates as for burden tests. We used the same five bins of gene constraint as used by the BHR authors. We then ran BHR on subsets of variants, *i.e.* selecting either synonymous, all missense, missense with PrimateAI-3D > 0.5, or LoF variants. We ran BHR separately for variants with MAF 0.1% -0.01%, and MAF < 0.01%, as recommended by the authors, and we summed the heritability for these disjoint groups within each variant category. We also ran BHR for missense variants using custom weights based on FlexRV burden tests significant at FDR 1%, with the idea that the burden weight will better reflect the likely effect size of each variant. However, since the FlexRV burden weight that gives the lowest p-value for a given gene is determined based on the same data as we use for BHR, we sought to mitigate overfitting. Instead of using the “best” burden weight function for each significant gene, we determined the burden weight function which gave a p-value equal to or higher (*i.e.* worse) than the multiple test-corrected p-value determined by CCT across burden weights. We used the weight function so determined for each significant gene to define the “custom_weights” parameter of BHR. For genes without a significant burden hit, we used the PrimateAI-3D score as the custom weight, since we did not have a burden weight. The heritability determined using the “multiple test-corrected” weight functions was lower than when using the best burden weight function for each gene, but was still higher than when using any fixed PrimateAI-3D threshold for missense variants or when using no threshold.

### Phenotype prediction

To compute polygenic scores (PGS) for traits based on rare variants, we first split 320,612 unrelated European ancestry individuals from UKB into 80%/20% train/test sets. We re-ran FlexRV on the training set samples and identified associations significant at FDR < 1%. We compared to the approach of Fiziev et al.^7^, which identifies the PrimateAI-3D score and MAF thresholds giving the strongest association, adjusting for multiple tests using permutations. Note that the procedure from Fiziev et al. selects a subset of missense and/or LoF variants in each gene, whereas FlexRV includes most or all variants while assigning a quantitative weight. We used the significant genes to train a new linear model per gene in the training set to predict phenotype values, which were IRNT-transformed residuals from covariate-adjusted phenotypes used in Fiziev et al. Rather than predicting phenotypes using the PrimateAI-3D score of variant carriers (and log(MAF)) as in Fiziev et al., we used the burden weight for the variant from the FlexRV burden weights which gave the lowest p-value in the training set. That is, we compared models trained using Y ∼ burden_weight + log(MAF) to models trained using Y ∼ score + log(MAF). Note that the burden weight already includes the impact of MAF, but we wanted to compare models with the same number of free parameters. We summed the predicted PGS for each individual across included genes. We used our model to predict phenotype values for samples in the test set and then evaluated the performance by considering the correlation R^2^ between predicted and observed phenotype values, using either PGS based on burden weights or by the method of Fiziev et al. (**Fig. 4b**). We also compared the enrichment of individuals having high/low PGS values (above/below 99%/1% quantiles of PGS, or 99.9%/0.1% quantiles) for having a phenotype value at the extreme of the distribution (sign-matched above/below a given z-score across individuals). These enrichments are plotted at different phenotype z-score thresholds (**Fig. 4c**) and showed a stronger enrichment of FlexRV-based PGS than Fiziev et al. PGS for identifying individuals with extreme phenotype values.

## Data availability

UK Biobank genetic and phenotype data are available to researchers with approved research applications (www.ukbiobank.ac.uk). Gene-trait associations significant at FDR 1% based on each method are provided in Supplementary Tables 2-3. S_het_ scores used herein were taken from supplementary table 1 of GeneBayes^35^. Full rare variant association test results will be made available at Zenodo upon publication.

## Code availability

The FlexRV package will be made available at https://github.com/Illumina/flexrv.

## Acknowledgements

This research has been conducted using the UK Biobank Resource under Application Number 33751. We gratefully acknowledge All of Us participants for their contributions and the National Institutes of Health’s All of Us Research Program, without whom this research would not have been possible. We thank Z. McCaw for providing helpful feedback.

